# Structure of the phosphoinositide 3-kinase p110γ-p101 complex reveals molecular mechanism of GPCR activation

**DOI:** 10.1101/2021.06.01.446612

**Authors:** Manoj K Rathinaswamy, Udit Dalwadi, Kaelin D Fleming, Carson Adams, Jordan TB Stariha, Els Pardon, Minkyung Baek, Oscar Vadas, Frank DiMaio, Jan Steyaert, Scott D Hansen, Calvin K Yip, John E Burke

**Author notes:** These authors contributed equally (Manoj K Rathinaswamy, Udit Dalwadi). To whom correspondence should be addressed: Calvin K Yip,; John E Burke.

## Abstract

The class IB phosphoinositide 3-kinase (PI3K), PI3Kγ, is a master regulator of immune cell function, and a promising drug target for both cancer and inflammatory diseases. Critical to PI3Kγ function is the association of the p110γ catalytic subunit to either a p101 or p84 regulatory subunit, which mediates activation by G-protein coupled receptors (GPCRs). Here, we report the cryo-EM structure of a heterodimeric PI3Kγ complex, p110γ-p101. This structure reveals a unique assembly of catalytic and regulatory subunits that is distinct from other class I PI3K complexes. p101 mediates activation through its Gβγ binding domain, recruiting the heterodimer to the membrane and allowing for engagement of a secondary Gβγ binding site in p110γ. Multiple oncogenic mutations mapped to these novel interfaces and enhanced Gβγ activation. A nanobody that specifically binds to the p101-Gβγ interface blocks activation providing a novel tool to study and target p110γ-p101-specific signaling events *in vivo*.

## Introduction

The class I phosphoinositide 3-kinase (PI3K) family of heterodimeric enzyme complexes are master regulators of numerous essential functions, including growth, survival, proliferation, and metabolism (*1*-*3*). Activation of PI3Ks downstream of cell-surface receptors leads to production of the lipid signal phosphatidylinositol 3,4,5, tris-phosphate (PIP_3_), which activates multiple downstream signaling pathways. The lipid kinase activity of class I PI3Ks is mediated by the p110 catalytic subunit of which there are four isoforms, split into class IA (p110α, p110β, p110δ) and class IB (p110γ) based on their association with distinct regulatory subunits. Class IB p110γ binds to either a p101 or p84 (also called p87) adaptor subunit (*4, 5*), which mediate activation by upstream stimuli.

The class IB p110 isoform p110γ, encoded by *PIK3CG* is a master regulator of immune cell function (*6*), chemotaxis (*7*), cytokine release (*8*), and reactive oxygen species generation (*9*), which are important processes for both the innate and adaptive immune systems. It is a key factor in multiple inflammatory diseases, including rheumatoid arthritis (*10*), atherosclerosis (*11*), Lupus (*12*), allergy (*8, 13, 14*), cardiovascular diseases (*15*), obesity related changes in metabolism (*16*), and pulmonary fibrosis (*17*). The immunomodulatory effects of PI3Kγ are drivers of pancreatic ductal adenocarcinoma (*18*) and targeting PI3Kγ in combination with checkpoint inhibitors has shown promise as an anti-cancer therapeutic (*19, 20*). Its ability to mediate multiple immune cell functions is controlled by its activation downstream of diverse cell surface receptors, including G-protein coupled receptors (GPCRs) (*21*), the IgE/Antigen receptor (*8*), receptor tyrosine kinases (*22*), and Toll-like receptors (TLRs) (*23, 24*).

Structural and biophysical analysis have provided initial insight into the regulation of p110γ and its activation by upstream stimuli. The structure of a p110γ fragment revealed a conserved molecular architecture shared by all p110 subunits composed of a Ras Binding Domain (RBD) which mediates activation by the small GTPase Ras (*25*), a C2 domain, an armadillo repeat helical domain, and an archetypal bi-lobal kinase domain, similar to protein kinases (*26*) (Fig. 1A). Sequence analysis suggested the presence of a ubiquitin-like domain at the N-terminus, possibly playing a similar function as the Adaptor Binding Domain (ABD) of class IA PI3Ks. Both p101 and p84 differ from the class IA PI3K regulatory subunits, as they are not essential for p110 stability and do not inhibit lipid kinase activity (*27*), but instead mediate activation by upstream stimuli. The molecular basis for why regulatory subunits differentially regulate class IA and IB PI3Ks has remained elusive.

**Figure 1.**
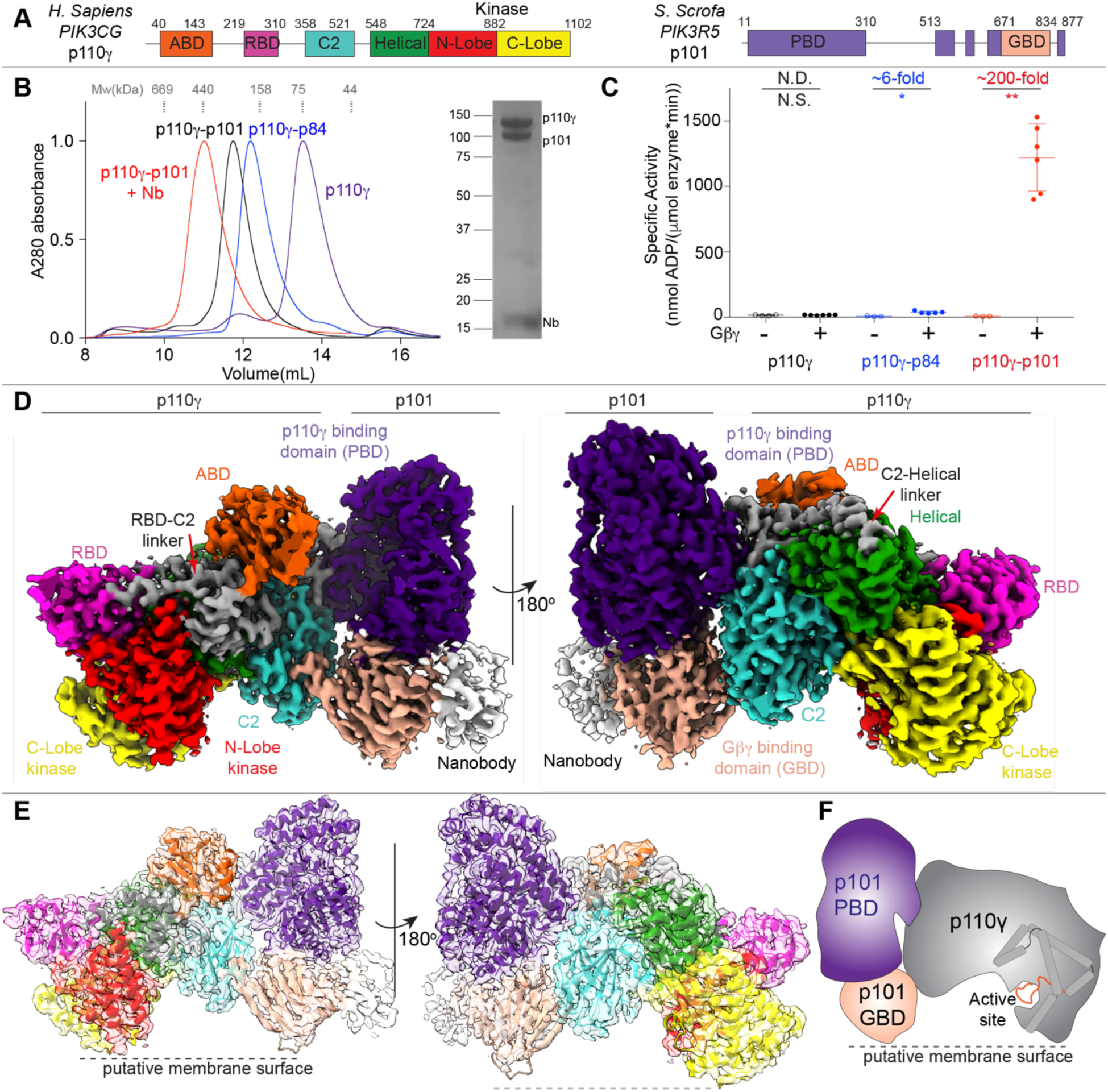
Cryo-EM structure of the p110γ p101 complex. **A.** Domain schematic of *H. Sapiens* p110γ and *S. Scrofa* p101 used in this study. **B.** Gel filtration elution profile of different p110γ complexes (i.e. apo or bound to p84, p101, and p101-NB1-PIK3R5). An SDS-PAGE image of the p110γ-p101-NB1-PIK3R5 complex is shown, with MW standards indicated. **C.** Lipid kinase activity assays of different p110γ complexes (concentration 30-3,000 nM) with and without lipidated Gβγ (1.5 μM concentration) using 5% PIP2 vesicles mimicking the plasma membrane (20% phosphatidylserine (PS), 50% phosphatidylethanolamine (PE), 10% Cholesterol, 10% phosphatidylcholine (PC), 5% sphingomyelin (SM) and 5% phosphatidylinositol-3,4,5-trisphosphate (PIP_2_)). The fold change upon Gβγ activation is indicated. Every replicate is plotted, with error shown as standard deviation (n = 3– 6). Two tailed p-values represented by the symbols as follows: **<0.001; *<0.02; N.S.>0.02. **D.** Density map of the p110γ-p101-NB1-PIK3R5 complex colored according to the schematic in panel **A**. **E.** Cartoon representation of the p110γ-p101 complex colored according to the schematic in panel **A**. **F.** Cartoon schematic of the p110γ-p101 complex.

The class I PI3Ks are frequently mis-regulated in multiple human diseases (*28*). This is most evident by the high frequency of hotspot somatic activating mutations found in *PIK3CA* (encodes for p110α) in multiple human cancers (*29, 30*). The other class I PI3K isoforms are also potentially involved in cancer, with overexpression of these p110 catalytic subunits leading to oncogenic transformation in cells (*31*). Both overexpression of *PIK3CG* and rare point mutations spanning the catalytic subunit have been identified in tumor biopsies (*32*-*34*). The mechanisms by which these mutations affect lipid kinase activity cannot be clearly explained by the existing structure of the catalytic subunit alone, highlighting the importance of understanding the molecular details of PI3Kγ regulatory complexes.

PI3Kγ is mainly activated downstream of GPCRs, where the presence of different adaptor subunits greatly modulates activation. The p84 and p101 subunits show distinct expression profiles, and alter PI3Kγ signaling responses to distinct upstream inputs (*35*). *In vivo,* the p110γ catalytic subunit alone is unable to be activated downstream of GPCRs and requires either the p101 or p84 regulatory subunits to respond to GPCRs (*36*). p101 and p84 play unique roles, with neutrophils lacking p84 having reduced reactive oxide species generation and neutrophils lacking p101 showing impaired migration(*37*). The distinct signaling responses were attributed to differential sensitivity of each of the PI3Kγ heterodimers to Gβγ subunits, with the p110γ-p101 complex being preferentially activated by Gβγ (*27, 38*) and p110γ-p84 requiring Ras binding for activation (*39*). To better understand how preferences for activating inputs translate into differences in function, there is a need for molecules that selectively inhibit one of the two p110γ complexes without affecting the other.

To decipher the molecular mechanism of how the p101 subunit regulates p110γ activation, we determined the structure of the p110γ-p101 complex using cryo-electron microscopy (cryo-EM). This structure reveals a novel binding interface between p101 and p110γ, which is completely distinct with the interface of class IA PI3K adaptors. Our structure also validates the presence of an ABD in p110γ similar to other class I PI3Ks, although with a unique orientation. Unlike class IA ABDs, the p110γ ABD does not directly bind p101, but instead orients the RBD-C2 linker for productive binding to this regulatory subunit. Intriguingly, oncogenic mutations found in p110γ localize at the interfaces of the ABD and p101. Hydrogen deuterium exchange mass spectrometry analysis revealed the altered dynamics of the p110γ-p101 interfaces upon mutation, leading to increased activation by Gβγ. The structure also showed that the Gβγ binding domain (GBD) in p101 contains a putative membrane binding surface that positions p110γ for catalysis. Single molecule fluorescence microscopy experiments indicated that the full activation of the p110γ-p101 complex requires the engagement of two Gβγ molecules to p101 and p110γ, respectively. Finally, a nanobody used in the cryo-EM analysis was found to be a potent inhibitor of GPCR activation of only p110γ-p101, with no effect on p110γ-p84. This nanobody could be used to decipher the complex specific roles of PI3Kγ in immune cell signaling, while also providing a novel potential therapeutic strategy for targeting unique PI3Kγ complexes.

## Results

### Structure of the p110γ-p101 complex

We purified full length human p110γ alone, as well as the p110γ-p84 and p110γ-p101 complexes (Fig. 1A+B). Gel filtration elution profiles of the p110γ-p84 and p110γ-p101 complexes confirmed their heterodimeric stoichiometry. Lipid kinase assays testing Gβγ activation revealed a ∼100-200-fold activation of p110γ-p101, a less potent ∼6-fold activation of p110γ-p84, and limited activation for p110γ alone, results that are consistent with previous work (*38*) (Fig. 1C). To delineate the molecular basis for how p101 protein controls the activation of p110γ we examined its architecture using an approach combining hydrogen deuterium exchange mass spectrometry (HDX-MS) and Cryo-EM.

We first conducted cryo-EM analysis of the p110γ-p101 complex. Although negative stain analysis revealed that purified p110γ-p101 was homogeneous (data not shown) and high-quality vitrified specimens from this relatively small-sized and asymmetric complex could be obtained, the region encompassing the p101 regulatory subunit was poorly resolved in our initial 3D reconstruction of p110γ-p101. This could be attributed to the highly dynamic nature of the C-terminal region of p101. To obtain a more “rigid” complex for cryo-EM analysis, we screened nanobodies targeting p110γ-p101 and found one that specifically stabilized the p101 C-terminal domain (NB1-PIK3R5, full details to be published in a separate manuscript). We purified the nanobody-bound p110γ-p101 complex and confirmed its 1:1:1 stoichiometry by gel filtration (Fig 1B). Using this sample, we were able to obtain a cryo-EM reconstruction of the ternary complex of nanobody-bound p110γ-p101 at 2.9 Å overall resolution from 320,179 particles (Table S2, Fig. S1-S3). The density map was of sufficient quality to allow for automated and manual building of the majority of the p110γ and p101 subunits (Fig. 1D-F). We were able to unambiguously fit available crystal structures of p110γ (144–1102) (*40*) into our map and build an additional 210 residues that constitute the ABD, the linkers connecting the RBD-C2 and C2-helical domains, and the kinase domain activation loop (Fig. S3C). The region with the lowest local resolution was the Gβγ binding domain (GBD) of p101, along with the bound nanobody. The GBD forms a beta sandwich structure composed of two sheets, and initial automated and manual model building only allowed for partial building of one of the two sheets. To build the remainder of the GBD we utilized a combination of Rosetta de novo modelling (*41*) and a trRosetta-guided protein folding method (*42, 43*) (Fig. S2). This allowed us to build a complete model of the structured regions of p101 (Fig. S4).

The p101 regulatory subunit structure (Fig. S4A-C) features a helical solenoid (11-149, 186-267) and a α/β barrel (150-185, 268-670, 867-877) that together we refer to as the p110γ binding domain (PBD), and a beta sandwich Gβγ binding domain (671-834, GBD). In addition to these motifs, there are four linker regions that were not resolved in the electron density map (311-512, 560-603, 623-650, and 835-866). Comparisons between p101 and p84 revealed that the structured regions of the PBD and GBD are partially conserved (28% identity and 47% similarity for the PBD, and 24% identity and 44% similarity for the GBD) (Fig. S4E). We analyzed our p101 model using the DALI server (*44*) and found multiple lipid kinases that shared a similar arrangement of an α/β barrel and β sandwich domain, including Diacylglycerol kinase, the phosphatidyl kinase YegS and Sphingosine kinase 1 (Fig. S4D). The catalytic residues at the active site were not conserved, suggesting that p101 has no kinase activity. However, comparing the previously identified lipid binding region of Sphingosine kinase 1 (*45*) with p110γ-p101 showed that the corresponding region of p101 is oriented towards the putative membrane interface (Fig. 1E+F), indicating a membrane binding site in p101.

### Molecular details of the p110γ-p101 interface

Our cryo-EM structure revealed that p101 engages p110γ in an extended interface resulting in buried surface area of ∼1202 Å^2^ (Fig. 2A+B, Fig. S5A+B). There are three specific regions of p110γ that bound to p101: the C2 domain, and the two linkers between the RBD-C2 and the C2-helical domains (Fig. 2A). These inter-domain linkers were not resolved in previous structures of p110γ alone and contain critical contact residues with p101. The primary binding interface on p101 is composed of the helices α5+6 and the intervening turn in the PBD which interacts with the C2 domain and linkers. The C2 domain also makes an additional interaction with the C-terminal proline of p101. These interfacial residues are strongly conserved across evolution (Fig. S4E). Intriguingly, we found that the PBD contact residues in p101 were also conserved with p84 (78% identical, 89% similar), revealing a likely shared mode of binding for both regulatory subunits.

**Figure 2.**
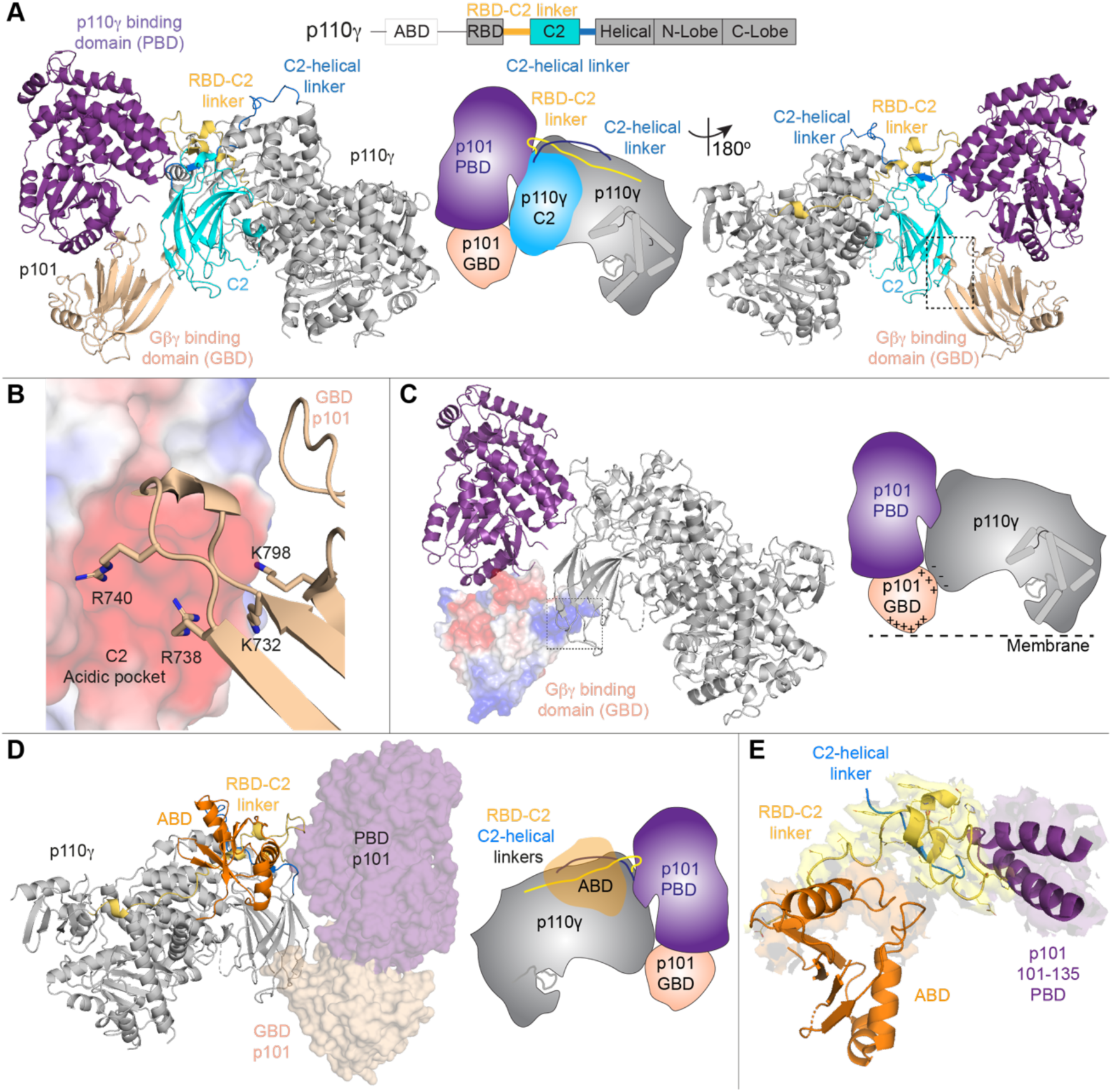
Structural basis of the p110γ-p101 binding interface. **A.** Cartoon representation of the p110γ-p101 complex, with p101 colored as in Figure 1, and p110γ colored according to the attached schematic, with p101 interacting regions (RBD-C2 linker, C2, and the C2-helical linker) indicated. Important features are shown in a cartoon schematic. **B.** Interaction between the GBD of p101 and the C2 domain of p110γ. The p110γ C2 domain is shown as an electrostatic surface with p101 shown as sticks. **C.** The electrostatic surface of the GBD of p101. A cartoon schematic highlighting potential electrostatic interactions between the GBD of p101 and the C2 domain of p110γ and membranes. **D.** The structure of p110γ-p101 complex, highlighting the orientation of the p110γ ABD, with p101 shown as a transparent surface. The different domains are colored as indicated according to the cartoon schematic. **E.** The ABD of p110γ coordinates the RBD-C2 linker of p110γ to interact with p101. The RBD-C2 linker, ABD, and the region of p101 that binds the RBD-C2 linker are shown in a cartoon representation. The electron density of the ABD interface, RBD-C2 linker and region of p101 that binds the RBD-C2 linker are visible.

A secondary contact site between p101 and p110γ is formed between two beta strands in the GBD of p101 (729–741) and the C2 domain of p110γ (Fig. S5B). This interface is formed by multiple electrostatic interactions between positively charged residues from the GBD of p101 and an anionic surface in the C2 domain of p110γ (Fig. 2B, Fig. S5B). This anionic surface in the C2 is absent in all other class I PI3Ks. Residues forming this contact site are conserved in the evolutionary history of both p101 and p110γ, but are only partially conserved in p84 (57% identical, 86% similar), suggesting that the dynamics of this contact may be altered between the two complexes. This unique interface could explain how a previously designed C2 binding antibody specifically inhibited GPCR activation of p110γ-p84 over p110γ-p101 (*46*).

To verify the contacts observed in the cryo-EM structure of p110γ-p101, and to compare to dynamics at the p110γ-p84 interfaces, we carried out HDX-MS experiments on p110γ alone and with the two regulatory subunits. Consistent with our previous work (*38, 47*), we found that with both p84 and p101, there was protection of large sections of the ABD and C2 domains along with the RBD-C2 and C2-helical linkers (Fig. S6A-C, Table S3). The same regions were protected in both complexes, however, the differences were larger in the presence of p101, indicating enhanced stability of the p101-bound complex. These differences in stability can be explained by the only partial conservation of the secondary interface residues between p101 and p84, in line with previous data (*35*).

Previous work showed that the interaction of the p110γ with its regulatory partners required the presence of the N-terminal ABD (*47*). Intriguingly, the ABD in p110γ does not directly interact with the regulatory subunit, but instead forms extensive contacts with the RBD-C2 linker (Fig. 2D, Fig. S5C+D) to orient the two linkers for binding the PBD of p101 (Fig. 2E, Fig. S5E). Consistent with this structural information, HDX-MS analysis of full length p110γ compared to a ΔABD construct (144–1102) showed clear protection of the RBD-C2 linker by the ABD (Fig. S6D).

### Comparison of regulatory subunit interactions in class IB to class IA PI3Ks

All class I PI3Ks bind to regulatory subunits, with the class IA PI3Ks binding to five different p85 regulatory subunits. p85 binding has three main effects: (1) it stabilizes the p110 catalytic subunit, (2) inhibits basal lipid kinase activity, and (3) allows activation downstream of pYXXM motifs (*48*). In contrast, association with class IB regulatory subunits neither stabilizes nor inhibits the p110γ catalytic subunit. To explain these differences in regulation, we compared the orientation of adaptor subunits and the ABD between class IA and class IB PI3Ks (Fig. 3A-D).

**Figure 3.**
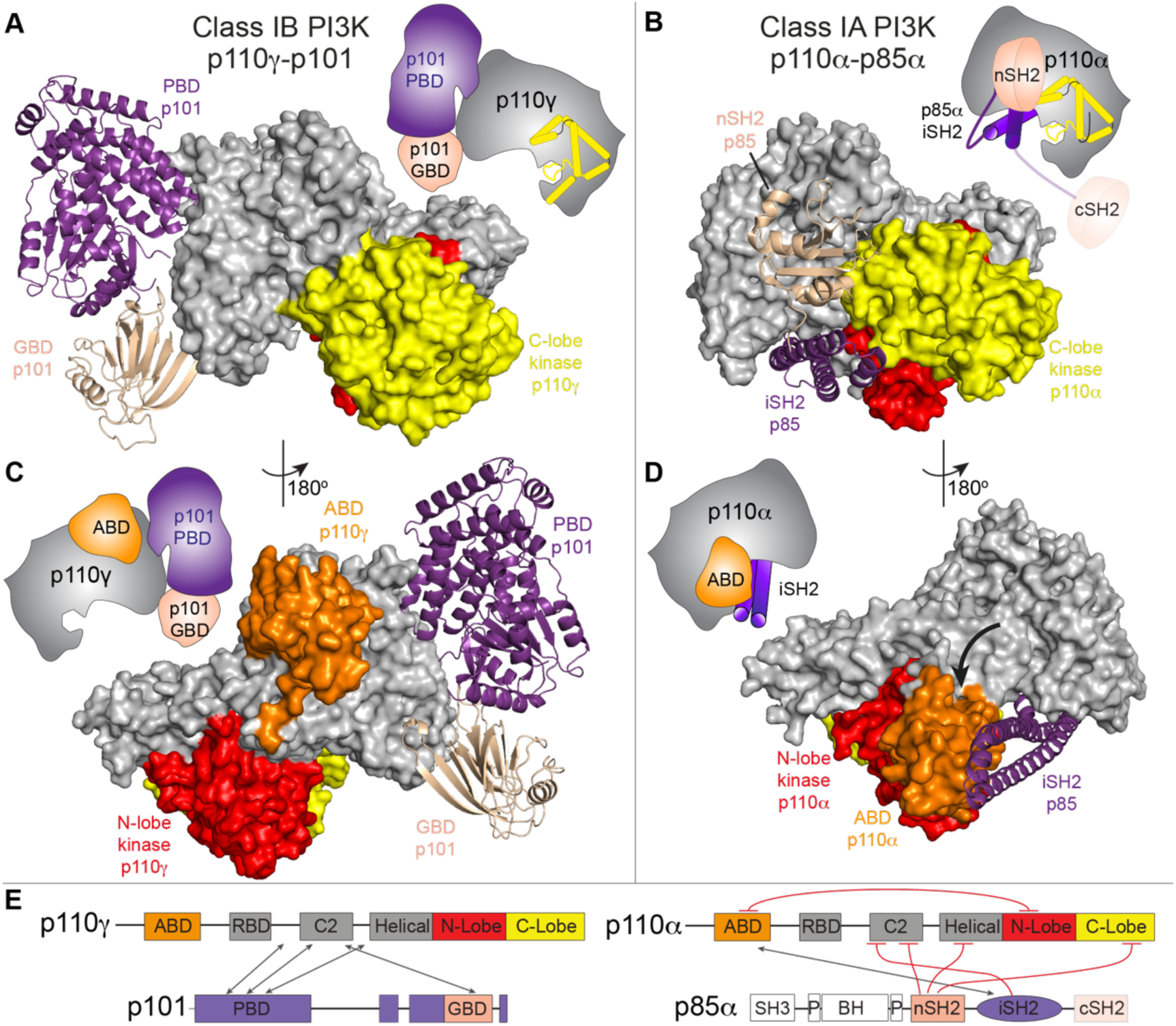
Class IA and IB PI3Ks form distinct interfaces with regulatory subunits and the ABD. **A.** The structure of p110γ-p101 complex, with p110γ shown as a surface and p101 shown as a ribbon, and the domains colored according to the cartoon schematic as indicated in panel **E**. **B.** The structure of p110α-p85α complex (PDB:4JPS), with p110α shown as a surface and the nSH2 and iSH2 domains of p85α shown as a ribbon, and the domains colored according to the cartoon schematic as indicated in panel **E**. **C.** The ABD of p110γ does not interact with either the regulatory subunit or kinase domain. The p110γ-p101 complex is shown as in panel **A**. **D.** The ABD of p110α interacts with both the regulatory subunit and kinase domain. The p110α-p85α complex is shown as in panel **B**. The altered orientation of the ABD compared to p110γ is indicated by the black arrow. (Cartoon schematics indicating the differences between class IA and class IB are shown for panels **A-D**). **E.** Domain schematic comparing the interactions between p110 catalytic and the p101 / p85 regulatory subunits. Inhibitory interactions are colored in red, with interacting regions indicated by the arrows.

The binding interface with regulatory subunits is completely distinct in class IA PI3Ks compared to class IB. While the C2 domain of class IA PI3Ks does interact with the iSH2 and nSH2 of p85 regulatory subunits, the interface is different from the one that binds the PBD and GBD of p101. Regulatory subunits in class IA PI3Ks make extensive inhibitory interactions with the C-lobe of the kinase domain, while no such contact is observed in the p110γ-p101 complex. Although ABDs of class IA and IB share a similar overall fold (Fig. S7A+B), there are extensive conformational differences in secondary structure elements, consistent with the limited sequence conservation (identity ranging from 13-16% for p110α, p110β, and p110δ). The ABD from class IA PI3Ks is required for forming a high affinity interaction with the iSH2 from p85, mediated by contacts between beta strands β1 and β2, and helix α3. This region is at the surface in the p110γ ABD, making no interactions with the rest of the p110γ subunit (Fig. S7C+D). For all class IA PI3Ks, the ABD forms an inhibitory contact with the N-lobe of the kinase domain, through its N-terminus and the β4-α3 loop. The ABD in p110γ is rotated 180° around the ABD-RBD linker allowing the β4-α3 loop to bind to the RBD-C2 linker. The residues in these regions are highly conserved in the evolution of p110γ, with no conservation with other class I PI3Ks (Fig. S7E+F). Overall, this comparison reveals why p110γ is not inhibited by p101 unlike class IA PI3Ks which are potently inhibited by extensive intra-and inter-subunit contacts with regulatory partners. (Fig. 3E).

### Rare oncogenic mutations cluster at the p101 and ABD interfaces in p110γ

The PI3K-Akt pathway is the most commonly activated pathway in human cancer, primarily driven by activating oncogenic mutations in the *PIK3CA* gene encoding p110α. The role of the p110γ isoform in cancer development has been ambiguous, with overexpression and rare somatic mutations of *PIK3CG* implicated in multiple cancers, including pancreatic, prostate, renal, and breast. To further explore if activating mutations potentially exist in *PIK3CG* we analyzed the Catalog Of Somatic Mutations In Cancer (COSMIC) (*49*), which showed mutations spanning the primary sequence of p110γ. Intriguingly, many of the most frequent mutations in p110γ mapped to interfaces with either the ABD or p101 (Fig. 4A).

**Figure 4.**
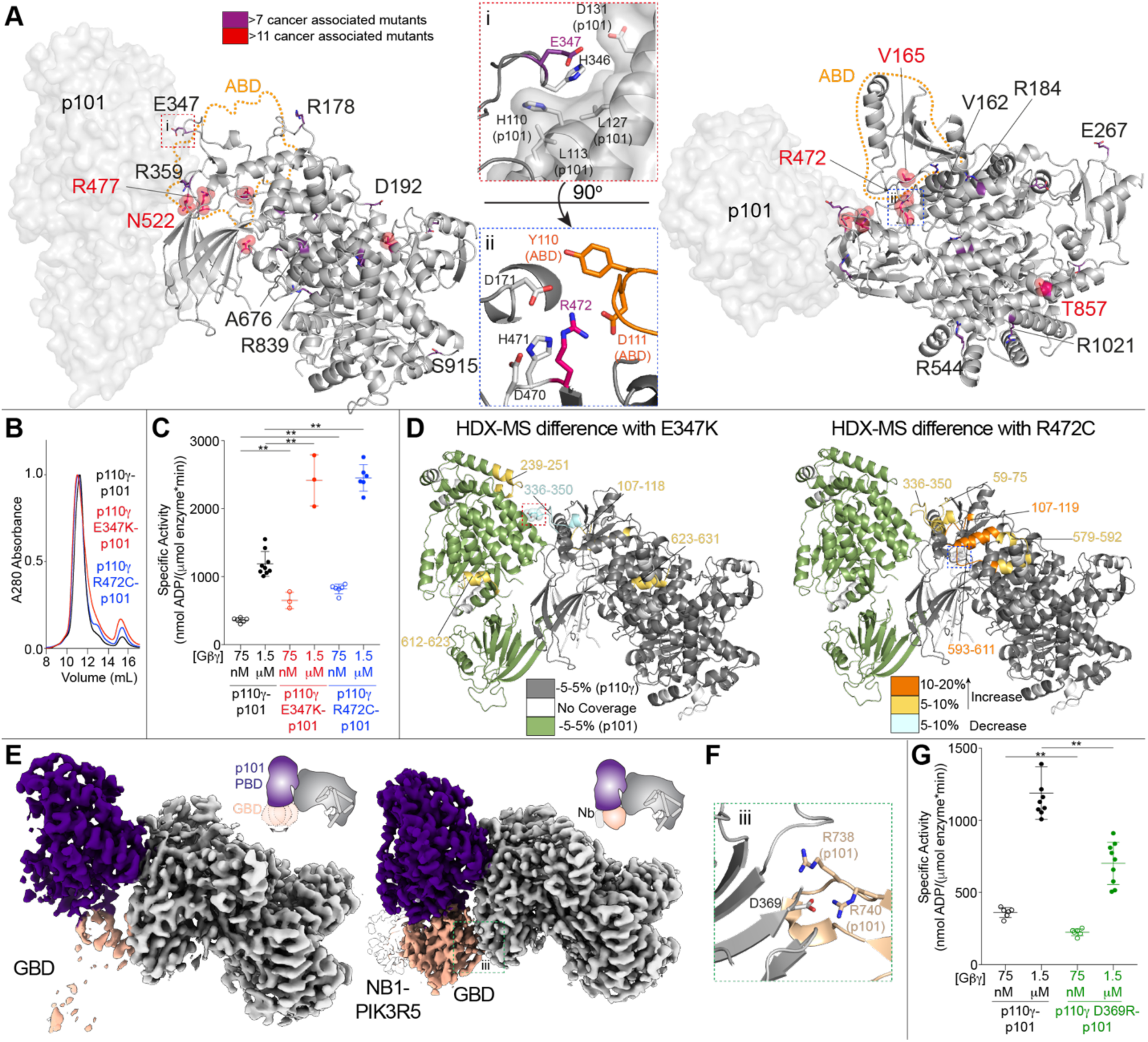
Disease-linked and engineered p110γ mutations at the interface with p101 and the ABD modulate Gβγ activation. **A.** Somatic mutations found in *PIK3CG* from the Catalogue of Somatic Mutations in Cancer Database (COSMIC) are indicated on the structure, with frequency indicated by the legend. Mutations found in more than 7 tumors are shown as sticks, with mutations found in more than 11 tumors shown as spheres. The orientation of residues around mutations located at the p110γ-p101 interface (i, E347) and ABD interface (ii, R472) are shown. **B.** Mutations do not disrupt the p110γ-p101 complex. Gel filtration elution profiles of complexes of p110γ (wild-type, E347, and R472) bound to p101. **C.** Mutations at the p110γ-p101 and ABD interface can lead to enhanced activation by Gβγ. Lipid kinase activity assays of different p110γ complexes (concentration 10-1,000 nM) with and without Gβγ (concentration indicated). **D.** Hydrogen deuterium exchange mass spectrometry (HDX-MS) revealed enhanced protein dynamics at p101 and ABD interfaces induced by E347K and R472 mutants. Peptides showing significant deuterium exchange differences (>5%,>0.4 kDa and p<0.01 in an unpaired two-tailed t-test) between p110γ-p101 complexes of wild-type and E347K (left) and wild-type and R472C (right) are colored on a cartoon model of p110γ-p101 according to the legend. **E.** The GBD is dynamic in solution, but is stabilized by nanobody (NB1-PIK3R5) binding. Electron density maps of p110γ-p101 alone (left) and p110γ-p101 bound to NB1-PK3R5 (right). **F.** Charged residues in p110γ-p101 mediate the interaction of the p110γ C2 domain to the p101 GBD. **G.** Mutation of the p110γ-C2 p101-GBD interface (p110γ D369R) leads to decreased activation by Gβγ. Biochemical assays in panels **C+G** were carried out with p110γ-p101 complexes (concentration 10-1,000 nM) and Gβγ (concentration indicated). 5% PIP_2_ membranes were made mimicking the plasma membrane. Every replicate is plotted, with error shown as standard deviation (n = 3–9). Two tailed p-values represented by the symbols as follows: **<0.001; *<0.02.

To identify the consequence of these interfacial mutations we purified mutant complexes of p101 with p110γ E347K (p101 interface) and p110γ R472C (ABD interface). Mutant complexes eluted from gel filtration similar to wild-type (Fig. 4B), demonstrating that they can still form heterodimers. Lipid kinase assays showed that the mutations resulted in a ∼2-3-fold increased activity at both saturating and sub-saturating amounts of Gβγ (Fig. 4C). To understand the mechanism of how these mutations lead to increased kinase activity, we carried out comparative HDX-MS experiments between the wild-type and mutant p110γ-p101 complexes. The E347K mutant caused increased dynamics at the p101 interface with p110γ, while the R472C mutant led to increased dynamics of the ABD, and the RBD-C2 linker (Fig. 4D, Fig. S8A+B). This suggests that altering the orientation of p101 and ABD relative to the rest of p110γ, may allow for increased access to membrane localized Gβγ subunits.

In addition to the disease-associated mutations at the primary interface between p110γ and p101, we also wanted to determine the role of the GBD-C2 interface in regulating Gβγ activation. This was motivated by analysis of the electron density maps between the free p110γ-p101 complex, and the p110γ-p101 complex with the NB1-PIK3R5 nanobody, which showed that the GBD is highly dynamic in the absence of the nanobody (Fig. 4E). We mutated a residue in the C2 domain (D369R) that interacts with R738 and R740 in the GBD (Fig. 4F). The p110γ D369R-p101 mutant eluted from gel filtration as a heterodimer, showing that this contact is not required for p101 binding. Lipid kinase activity assays revealed that this mutation led to a ∼2-fold decrease in activation by Gβγ subunits, indicating that the secondary p110γ-p101 interface is crucial in mediating full activation (Fig. 4G).

### The Gβγ binding domain is critical for membrane binding and Gβγ activation

To decipher how p101 mediates activation by Gβγ, we have previously examined the dynamic consequences of p110γ-p101 binding to membranes, and membrane localized Gβγ subunits using HDX-MS (*38*). Our current model of p110γ-p101 allowed us to better understand these data in the context of the full complex. Analysis of the HDX differences upon membrane binding indicated the presence of a membrane binding region in the GBD and protection at the secondary interface between p110γ and p101 (Fig. 5A, S8C+E). Upon binding to Gβγ subunits, many of these same regions showed greatly decreased exchange indicating enhanced membrane recruitment. Additionally, the Gβγ binding sites (p101 GBD and p110γ helical domain) showed decreases in exchange (Fig. 5B, S8D+E). Combined with our observations of the GBD flexibility in cryo-EM and activity assays with the p110γ D369R mutant, this HDX-MS data indicated that the GBD forms the secondary interface upon membrane binding, potentially explaining the inability of non-lipidated soluble Gβγ (C68S, Gγ) to interact with p110γ-p101.

**Figure 5.**
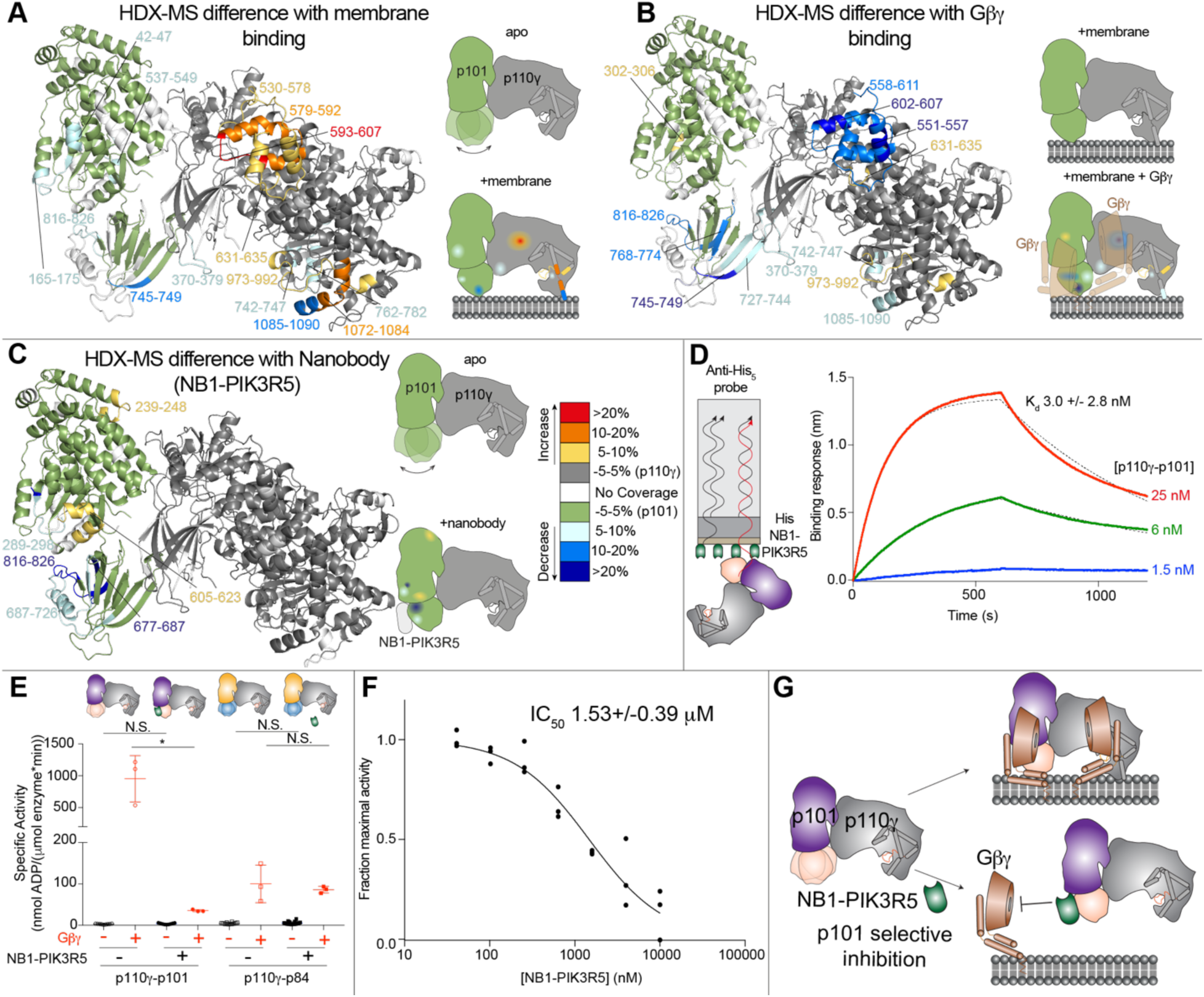
Full activation of p110γ by lipidated Gβγ requires the GBD domain of p101 and the GBD-Gβγ interface can be disrupted by a p101 specific nanobody. **A.** HDX-MS revealed that interaction of p110γ-p101 with membranes leads to altered protein dynamics in both the p110γ and p101 subunits, with stabilization of the GBD of p101. For panels **A-C**, peptides showing significant deuterium exchange differences (>5%,>0.4 kDa and p<0.01 in an unpaired two-tailed t-test) between conditions are colored on a cartoon model of p110γ-p101 according to the legend in panel B. A cartoon schematic is shown indicating the two conditions compared using HDX-MS. **B.** HDX-MS revealed that interaction of p110γ-p101 with lipidated Gβγ subunits stabilizes the GBD and C2-helical/helical domain of p110γ. HDX-MS data from panels **A+B** are reproduced with permission from (*38*). **C.** HDX-MS revealed that interaction of p110γ-p101 with NB1-PIK3R5 protects the same surface of GBD that is stabilized upon binding Gβγ on membranes. **D.** Biolayer interferometry (BLI) analysis of the binding of the immobilized NB1-PIK3R5 nanobody to p110γ-p101. **E.** The NB1-PIK3R5 nanobody specifically inhibits only the p110γ-p101 complex from GPCR activation, while not affecting the p110γ-p84 complex. Biochemical assays were carried out with p110γ-p101 (50-3,000 nM) and p110γ-p84 (1,500-3,000 nM) using plasma membrane mimic vesicles with and without NB1-PK3R5 (6 μM). Lipidated Gβγ was present at 1.5 μM concentration. **F.** IC50 measurement of p110γ-p101 inhibition using varying concentrations of the NB1-PIK3R5 nanobody in the presence of 600 nM Gβγ. For panels **E**+**F** every replicate is plotted, with error shown as standard deviation (n = 3–6). Two tailed p-values represented by the symbols as follows: **<0.001; *<0.02. N.S.>0.02. **G.** Model of the inhibition of GPCR activation of the p110γ-p101 complex by the NB1-PIK3R5 nanobody.

### A p101 binding nanobody prevents activation by Gβγ subunits

HDX-MS analysis of the NB1-PIK3R5 nanobody used in cryo-EM experiments of p110γ-p101(Fig. 1D, 5C, Fig. S8F) showed that it binds with high affinity (∼3 nM) to the identified Gβγ binding site in the GBD (Fig. 5D). This suggested that this nanobody might be useful in specifically disrupting Gβγ activation of the p110γ-p101 complex. We utilized lipid kinase assays with both p110γ-p101 and p110γ-p84 to study the effects of NB1-PIK3R5 on GPCR activation. The NB1-PIK3R5 nanobody at 6 μM led to a ∼50-100-fold reduction in Gβγ activation for p110γ-p101, with no effect on p110γ-p84 activation (Fig. 5E). The nanobody was capable of potently inhibiting the p110γ-p101 complex (IC50 ∼1.5 μM) at super-physiological levels of Gβγ (600 nM) (Fig. 5F), thereby providing a novel tool that may aid in deciphering the exact roles of the p110γ-p101 complex in cells/tissues (Fig. 5G), and in designing complex-specific therapeutic strategies.

### Defining the molecular basis of how Gβγ subunits activate the p110γ-p101 complex

Structural analysis of the Gβγ binding sites in p110γ and p101 from HDX-MS indicated that these regions are separated by ∼50 Å, which is greater than the ∼40 Å diameter of the Gβγ propeller domain. This suggested that activation of p110γ-p101 is potentially mediated by interactions with two membrane anchored Gβγ molecules. To gain new insight about the mechanism of Gβγ dependent activation of p110γ-p101 we performed single molecule Total Internal Reflection Fluorescence (TIRF) Microscopy experiments on supported lipid bilayers (SLBs) (Fig. 6A, S9). Experiments were carried out using fluorescently tagged proteins (DY647-p110γ, DY647-p110γ-p101, and Alexa488-SNAP-Gβγ) to track membrane binding. For these experiments, we flowed farnesylated Alexa488-SNAP-Gβγ over a SLB, leading to passive insertion into the membrane (t_1/2_ ∼8 min; Fig. S9). Single molecule dwell time measurements of fluorescently tagged DY647-p110γ and DY647-p101-p110γ revealed no appreciable membrane binding in the absence of Gβγ (Fig. 6B). In the presence of membrane anchored Gβγ, we observed an increased binding frequency of DY647-p110γ and transient dwell times that lasted 10-100 ms (Figure 6B+C). By contrast, DY647-p101-p110γ bound strongly to membrane anchored Gβγ and exhibited single molecule dwell times that lasted several seconds (Figure 6B+D, Table S4).

**Figure 6.**
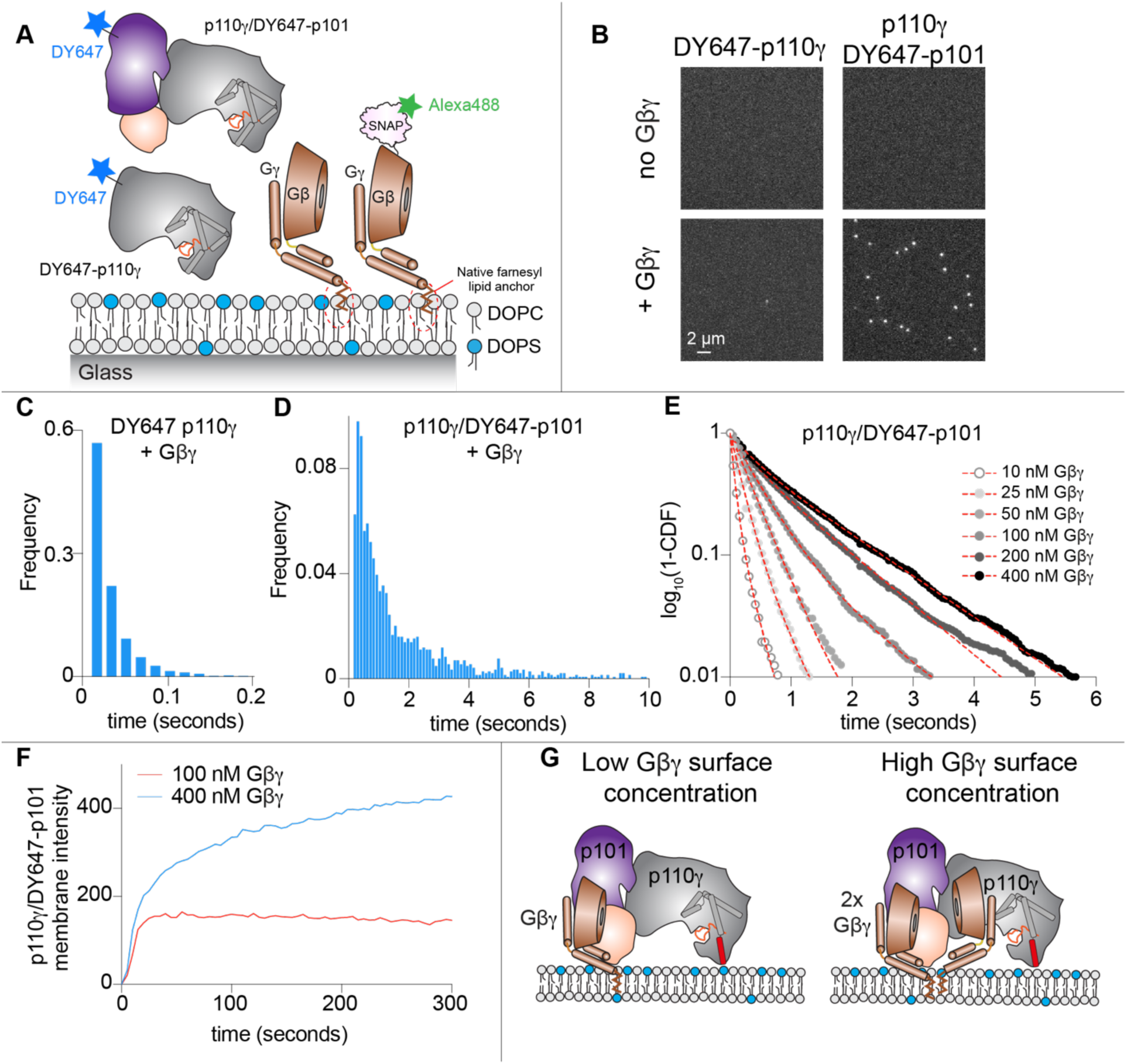
Single molecule characterization of p110γ-p101 reveals both subunits can engage membrane anchored Gβγ. **A.** Schematic showing proteins examined using the single molecule fluorescence approach. Experiments measured the association of fluorescently tagged proteins (Alexa488-SNAP-Gβγ, DY647**-**p110γ, and DY647-p101-p110γ to a supported lipid bilayer. **B.** Membrane association of DY647**-**p110γ or DY647-p101-p110γ requires membrane anchored Gβγ. Single molecule localization measurements were measured in the presence of either 100 pM DY647**-**p110γ or 10 pM DY647-p101-p110γ. **C-D.** Single molecule dwell time distributions of DY647**-**p110γ or DY647-p101-p110γ, measured in the presence of membrane anchored Gβγ. DY647**-**p110γ transiently associates with membrane anchored Gβγ (**C** τ_1_= 22 ms, n=2832 events). DY647-p101-p110γ binds strongly to membrane anchored Gβγ (τ_1_= 0.334 sec (31%), τ_2_= 1.31 sec (69%), n=3996 events). **E.** Gβγ membrane density dependent changes in the membrane binding behavior of DY647-p101-p110γ. Concentration of Gβγ represents the solution concentration. **F.** DY647-p101-p110γ absorption kinetics at different Gβγ membrane densities **G.** Model of p110γ-p101 recruitment to Gβγ subunits at both low and high membrane densities.

Since both subunits in the p110γ-p101 complex contain Gβγ binding interfaces we hypothesized that the single molecule dwell times of DY647-p101-p110γ would strongly depend on the concentration of membrane anchored Gβγ. When we titrated the concentration of Gβγ, we observed a density dependent switch in p110γ-p101 membrane binding behavior (Fig. 6E, Table S4). In the presence of low Gβγ concentration (i.e. ≤ 100 nM), the dwell time distribution of DY647-p101-p110γ was best described by a single exponential decay curve with dwell times ranging from 100-400 ms (Table S4). In contrast, we observed longer lived membrane binding interactions when our measurements were performed using more than 100 nM Gβγ. Under these conditions, the dwell time distribution shifted from a single to double exponential decay curve (Table S4).

Consistent with the concentration of Gβγ modulating the dwell time of DY647-p101-p110γ, we also observed changes in the bulk membrane absorption kinetics of p110γ-p101 (Fig. 6F). In the presence of low Gβγ concentrations (100 nM), DY647-p101-p110γ rapidly associated with the membrane and reached an equilibrium within ∼30 seconds (Fig. 6F). This is the expected kinetic profile for a simple biomolecular interaction between two proteins. In contrast, the membrane absorption kinetics of DY647-p101-p110γ was biphasic in the presence of 400 nM Gβγ (Fig. 6F). Under these conditions, DY647-p101-p110γ association kinetics are described by rapid binding to the first Gβγ, followed by slow engagement with a second Gβγ (Fig. 6F). This type of biphasic membrane absorption is similar to how BTK reportedly interacts with two PI(3,4,5)P_3_ lipids (*50*). In summary, our TIRF microscopy measurements show that the p110γ-p101 can engage up to two Gβγ molecules depending on the level of GPCR activation (Fig. 6G).

## Discussion

Understanding how p110γ activity is regulated by p84 or p101 regulatory subunits has been critical in deciphering physiological roles (*13, 36, 37*), and will be important in effective PI3K therapeutic design. The class IB p110γ catalytic subunit is a key regulator of immune cell signaling and is a therapeutic target for inflammatory diseases (*10, 51*) and immunomodulatory cancer treatment (*19, 20*). Here we report the architecture of the p110γ-p101 complex and a new mechanism of how it can be activated during GPCR signaling.

Our cryo-EM structure of p110γ-p101 reveals important differences in the assembly of catalytic and regulatory subunits between class IA and class IB PI3Ks, and provides novel insight into PI3K regulation. Previous X-ray crystallographic studies of a p110γ fragment revealed its domain organization and the molecular basis for interaction with the upstream activator Ras (*25, 26, 40*). Our cryo-EM structure showed an evolutionarily well-conserved binding surface between the regulatory p101 subunit and p110γ. The p110γ ABD does not directly bind p101, but instead coordinates the p101 interaction site on the RBD-C2 linker. This unique architecture is distinct from the ABD in class IA PI3Ks, which mediates direct contacts with the iSH2 of p85 and forms an inhibitory interface with the N-lobe of the kinase domain (*52, 53*).

The architecture of the p110γ-p101 complex reveals how the GBD of p101 orients the kinase domain of p110γ towards the membrane upon Gβγ binding. The p101 protein was identified as a key regulator of the activation of p110γ by GPCRs (*5, 54*), however, defining the full details of this mechanism was hampered by a paucity of structural information. Complicating structural analysis of p101 is the lack of homologous proteins, with p84 being the only protein with greater than 20% sequence identity in the human proteome. Our p101 structural model validated and defined at high resolution the presence of a Gβγ binding surface on the GBD of p101 (*38*). The structural similarity of the α/β barrel and the GBD of p101 with the lipid binding surfaces of the lipid kinases diacylglycerol kinase and sphingosine kinase (*45*) support the idea that the GBD participates in membrane binding upon Gβγ activation. Cellular studies revealed that Gβγ was able to activate membrane localized p110γ, suggesting that Gβγ may orient p110γ in a catalytically competent state (*55*). Our structure reveals how the GBD participates in membrane and Gβγ binding, which orients the kinase domain for catalysis. Once recruited to the membrane, the p110γ-p101 complex can engage a second Gβγ binding site located on the helical domain of p110γ, leading to full activation. Further structural analysis of the GBD bound to Gβγ will be required to narrow down the exact molecular details of GPCR activation.

The ability of p110γ to generate discrete PIP_3_ signals upon activation by a unique set of upstream stimuli is critical to their role in immune cells (*1*) and is ultimately controlled by the p101 and p84 regulatory subunits. This has been highlighted by responses in neutrophils and mast cells, where p101 complexes mediate cell migration, and p84 complexes mediate reactive oxide production and degranulation (*13, 37*). Various stimuli have been identified that can activate p110γ, including G-protein coupled receptors (*27*), Ras (*25*), Toll like receptors (TLR, mediated by Rab8 activation) (*24, 56*), and the IgE antigen receptor (partially mediated by protein kinase C phosphorylation of p110γ) (*47*). The p101 and p84 regulatory subunits confer the ability to be preferentially stimulated by a specific subset of these stimuli. The p110γ-p84 complex is uniquely sensitive to Ras activation (*39*), and is less responsive to signals downstream of GPCRs in comparison to the p101 complex (*36*). Our TIRF microscopy data reveals how p110γ-p101 is uniquely situated to generate varying PIP_3_ responses depending on the different Gβγ membrane surface densities (Fig. 7A). The GBD of p101 uniquely responds to low Gβγ surface densities and allows for distinct p110γ-p101 stimulated PIP_3_ responses compared to p110γ-p84. In combination with the membrane localized activators Ras and Rab8 this allows for a multi-faceted set of PIP_3_ responses generated by p110γ-p101.

**Figure 7.**
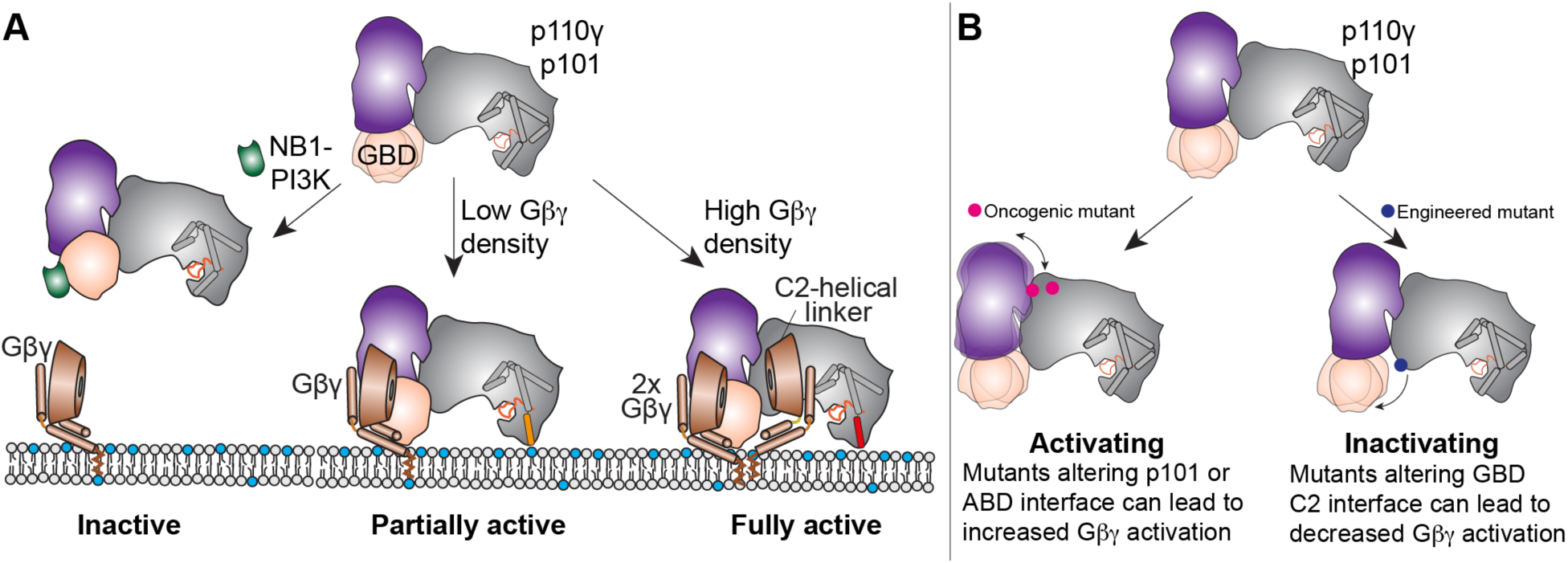
Model for regulation of p110γ-p101 activation by Gβγ membrane density, and modulation by nanobodies and disease-linked mutations. **A.** Schematic of how Gβγ subunits can lead to p110γ-p101 activation at different Gβγ surface densities, and how this can be disrupted by the NB1-PIK3R5 nanobody. **B.** Schematic of how mutations at the p101 and ABD interfaces in p110γ can lead to enhanced Gβγ activation, and how disruption of the GBD-C2 interface can lead to decreased Gβγ activation.

Activating mutations in the class I PI3K pathway are the most frequent alterations in human cancer (*30*), with this primarily driven by oncogenic mutations of p110α (*29, 57*). p110γ is often overexpressed in cancer, specifically in pancreatic ductal adenocarcinoma (*18, 33, 58*). Supporting this role of increased expression of p110γ in cancer is the knockdown of a microRNA targeting *PIK3CG* in patients that promotes metastasis in triple negative breast cancer (*32*). Tumor associated mutations in p110γ are rare compared to p110α (1739 for *PIK3CG* compared to 17,359 for *PIK3CA,* from the COSMIC database as of April 2021 (*49*)). However, multiple studies have found an association of somatic mutations in *PIK3CG* with cancer development and progression (*59*-*61*). Whether these mutants function within the tumor or the surrounding immune environment remains to be confirmed. Activating oncogenic mutants in the regulatory motif of the kinase domain of p110γ (R1021C) have been identified (*62*), with bi-allelic inactivating mutations involving the same site (R1021P, R982 frameshift) causing primary immunodeficiencies (*63*). We found oncogenic mutations clustered at ABD and p101 interfaces in *PIK3CG.* These mutants led to increased kinase activity upon Gβγ stimulation, which is explained by the altered interaction between p110γ and p101 as observed by HDX-MS. This may lead to a reorientation of the GBD allowing for increased binding to membranes or lipidated Gβγ (Fig. 7B). Further analysis of the effect of these mutations on membrane recruitment, and their effects in cells and model organisms, will be required to understand their complete mechanism of regulation.

The association of p110γ in human disease has driven intense interest in the generation of PI3K selective small molecule inhibitors, however, severe side-effects have limited their efficacy, particularly for pan-PI3K inhibitors (*64*). Multiple p110γ isoform selective inhibitors are currently in clinical trials for cancer, and are in development for COPD, and inflammatory disease. Inhibition of p110γ has also been found to improve anti-tumor properties of CAR T-cells (*65*). Regulatory subunits are differentially involved in the onset and progression of p110γ associated diseases. Upregulation of p110γ-p101 is involved in congestive heart failure (*66*), while p110γ-p84 plays a protective role by maintaining cardiac contractility (*15*). The p110γ-p101 complex could also be involved in TLR9-induced inflammation (*67*) due to its sensitivity to Rab8 activation downstream of TLRs (*23*). In pancreatic cancer models, targeting p110γ is protective in cancer development (*33*), however, its applicability is limited by hepatotoxicity. Therefore, targeting p110γ in these disease states could benefit from specifically inhibiting p110γ-p101 signaling. We have identified the structural basis for how the NB1-PIK3R5 nanobody can selectively inhibit Gβγ activation of the p110γ-p101 complex, which can be used to determine potential advantages of p110γ-p101 specific inhibition in p110γ-linked diseases, and may allow for design of novel therapeutic strategies.

Collectively, our detailed biochemical and structural analysis of the p110γ-p101 complex provides unique insight into the assembly and regulation of PI3Kγ complexes. This work provides a framework for the design of selective modulators outside of the ATP binding pocket, which will be useful to decipher PI3Kγ signaling roles and for the generation of potential therapeutics in inflammatory diseases and cancer.

## Materials and Methods

(full resources for all experiments in Table S1)

### Expression and purification of nanobody

Nanobody NB1-PIK3R5 with a C-terminal 6X His tag was expressed from a pMESy4 vector in the periplasm of WK6 E.coli. A 1L culture was grown to OD600 of 0.7 in Terrific Broth containing 0.1% glucose and 2mM MgCl2 in the presence of 100 μg/mL ampicillin and was induced with 0.5 mM isopropyl-β-D-thiogalactoside (IPTG). Cells were harvested the following day by centrifuging at 2500 RCF (Eppendorf Centrifuge 5810 R) and the pellet was snap-frozen in liquid nitrogen. The frozen pellet was resuspended in 15 mL of buffer containing 200 mM Tris pH 8.0, 0.5mM ethylenediaminetetraacetic acid (EDTA) and 500 mM Sucrose and was mixed for 1 hour at 4°C. To this mixture, 30 mL of resuspension buffer diluted four times in water was added and mixed for 45 minutes at 4°C to induce osmotic shock. The lysate was clarified by centrifuging at 14,000 rpm for 30 minutes (Beckman Coulter JA-20 rotor). Imidazole was added to the supernatant to final concentration of 10mM loaded onto a 5 mL HisTrap™ FF crude column (GE Healthcare) equilibrated in NiNTA A buffer (20 mM Tris pH 8.0, 100 mM NaCl, 20 mM imidazole pH 8.0, 5% (v/v) glycerol, 2 mM β-mercaptoethanol (βME)). The column was washed with high salt NiNTA A buffer (20 mM Tris pH 8.0, 1 M NaCl, 20 mM imidazole pH 8.0, 5% (v/v) glycerol, 2 mM βME), NiNTA A buffer, 6% NiNTA B buffer (20 mM Tris pH 8.0, 100 mM NaCl, 250 mM imidazole pH 8.0, 5% (v/v) glycerol, 2 mM βME) and the protein was eluted with 100% NiNTA B. The eluent was concentrated in a 10,000 MWCO Amicon Concentrator (Millipore) to <1 mL and injected onto a Superdex™ 75 10/300 GL Increase size-exclusion column (GE Healthcare) equilibrated in gel filtration buffer (20mM Tris pH 8.5, 100 mM NaCl, 50 mM Ammonium Sulfate and 0.5 mM tris(2-carboxyethyl) phosphine (TCEP)). Following size exclusion, the protein was concentrated, frozen and stored at −80°C.

### Plasmid Generation for PI3Kγ constructs

PI3Kγ constructs without the regulatory subunit (p110γ full length and p110γ 144-1102) were encoded in a pACEBac vector while the complexes were expressed from MutliBac (WT) or biGBac (mutants) vectors. For purification, a 10X histidine tag, a 2X-strep tag and a Tobacco Etch Virus protease cleavage site were cloned to the N-terminus of the regulatory subunits for the complex and to p110γ for constructs without regulatory subunits. All mutations were made in pLib vectors encoding p110γ using site-directed mutagenesis according to published commercial protocols (QuickChange Site-Directed Mutagenesis, Novagen). Oligonucleotides spanning the region of interest containing altered nucleotides were used in PCR reactions (Q5 High-Fidelity 2X MasterMix, New England Biosciences #M0492L) and the resulting reaction mixture was transformed into XL10 E.coli. Single colonies were grown overnight and purified using QIAprep Spin Miniprep Kit (Qiagen #27104). Plasmid identity was confirmed by sequencing.

### Virus Generation and Amplification

The plasmids encoding genes for insect cell expression were transformed into DH10MultiBac cells (MultiBac, Geneva Biotech) containing the baculovirus viral genome (bacmid) and a helper plasmid expressing transposase to transpose the expression cassette harbouring the gene of interest into the baculovirus genome. Bacmids with successful incorporation of the expression cassette into the bacmid were identified by blue-white screening and were purified from a single white colony using a standard isopropanol-ethanol extraction method. Briefly, colonies were grown overnight (16 hours) in 3-5 mL 2xYT (BioBasic #SD7019). Cells were pelleted by centrifugation and the pellet was resuspended in 300 μL P1 Buffer (50 mM Tris-HCl, pH 8.0, 10 mM EDTA, 100 mg/mL RNase A), chemically lysed by the addition of 300 μL Buffer P2 (1% sodium dodecyl sulfate (SDS) (W/V), 200 mM NaOH), and the lysis reaction was neutralized by addition of 400 μL Buffer N3 (3.0 M potassium acetate, pH 5.5). Following centrifugation at 21130 RCF and 4 °C (Rotor #5424 R), the supernatant was separated and mixed with 800 μL isopropanol to precipitate the DNA out of solution. Further centrifugation at the same temperature and speed pelleted the Bacmid DNA, which was then washed with 500 μL 70% Ethanol three times. The Bacmid DNA pellet was then dried for 1 minute and re-suspended in 50 μL Buffer EB (10 mM Tris-Cl, pH 8.5; All buffers from QIAprep Spin Miniprep Kit, Qiagen #27104). Purified bacmid was then transfected into Sf9 cells. 2 mL of Sf9 cells between 0.3-0.5X10^6^ cells/mL were aliquoted into the wells of a 6-well plate and allowed to attach, creating a monolayer of cells at ∼70-80% confluency. Transfection reactions were prepared by the addition of 2-10 μg of bacmid DNA to 100 μL 1xPBS and 12 μL polyethyleneimine (PEI) at 1 mg/mL (Polyethyleneimine ‘‘Max’’ MW 40.000, Polysciences #24765, USA) to 100 μL 1xPBS. The bacmid-PBS and the PEI-PBS solutions were mixed together, and the reaction occurred for 20-30 minutes before addition drop-by-drop to an Sf9 monolayer containing well. Transfections were allowed to proceed for 5-7 days before harvesting virus containing supernatant as a P1 viral stock.

Viral stocks were amplified by adding P1 viral stock to suspension Sf9 cells between 1-2×10^6^ cells/mL at a 2/100 volume ratio. This amplification produces a P2 stage viral stock that can be used in final protein expression. The amplification proceeded for 4-5 days before harvesting, with cell shaking at 120 RPM in a 27°C shaker (New Brunswick). Harvesting of P2 viral stocks was carried out by centrifuging cell suspensions in 50 mL Falcon tubes at 2281 RCF (Beckman GS-15), collecting the supernatant in a fresh sterile tube, and adding 5-10% inactivated foetal bovine serum (FBS; VWR Canada #97068-085).

### Expression and purification of PI3Kγ constructs

The PI3Kγ complexes (Human p110γ-porcine p101 WT/mutants and Human p110γ-mouse p84 WT) were expressed in Sf9 insect cells using the baculovirus expression system. Following 55 hours of expression, cells were harvested by centrifuging at 1680 RCF (Eppendorf Centrifuge 5810 R) and the pellets were snap-frozen in liquid nitrogen. Constructs without the regulatory subunit (Human p110γ full length and Human p110γ 144-1102) were expressed in insect cells for 55 hours from a pACEBac vector. Both the monomer and the complex were purified identically through a combination of nickel affinity, streptavidin affinity and size exclusion chromatographic techniques.

Frozen insect cell pellets were resuspended in lysis buffer (20 mM Tris pH 8.0, 100 mM NaCl, 10 mM imidazole pH 8.0, 5% glycerol (v/v), 2 mM βME), protease inhibitor (Protease Inhibitor Cocktail Set III, Sigma)) and sonicated for 2 minutes (15s on, 15s off, level 4.0, Misonix sonicator 3000). Triton-X was added to the lysate to a final concentration of 0.1% and clarified by spinning at 15,000 RCF for 45 minutes (Beckman Coulter JA-20 rotor). The supernatant was loaded onto a 5 mL HisTrap™ FF crude column (GE Healthcare) equilibrated in NiNTA A buffer (20 mM Tris pH 8.0, 100 mM NaCl, 20 mM imidazole pH 8.0, 5% (v/v) glycerol, 2 mM βME). The column was washed with high salt NiNTA A buffer (20 mM Tris pH 8.0, 1 M NaCl, 20 mM imidazole pH 8.0, 5% (v/v) glycerol, 2 mM βME), NiNTA A buffer, 6% NiNTA B buffer (20 mM Tris pH 8.0, 100 mM NaCl, 250 mM imidazole pH 8.0, 5% (v/v) glycerol, 2 mM βME) and the protein was eluted with 100% NiNTA B. The eluent was loaded onto a 5 mL StrepTrap™ HP column (GE Healthcare) equilibrated in gel filtration buffer (20mM Tris pH 8.5, 100 mM NaCl, 50 mM Ammonium Sulfate and 0.5 mM TCEP). The column was washed with the same buffer and loaded with tobacco etch virus protease. After cleavage on the column overnight, the protein was eluted in gel filtration buffer. For the complex with nanobody, the eluted protein was incubated with two-fold molar excess of purified nanobody on ice for 15 minutes. The protein was concentrated in a 50,000 MWCO Amicon Concentrator (Millipore) to <1 mL and injected onto a Superdex™ 200 10/300 GL Increase size-exclusion column (GE Healthcare) equilibrated in gel filtration buffer. After size exclusion, the protein was concentrated, aliquoted, frozen and stored at −80°C.

### Cryo-EM Sample Preparation and Data Collection

C-Flat 2/2-T 300 mesh grids were glow discharged for 25s at 15mA using a Pelco easiGlow glow-discharger. 3μL of purified p110γ-p101 complex with or without bound nanobody was then applied to the grids at a concentration of 0.45 mg/ml. Grids were then prepared using a Vitrobot Mark IV (Thermo Fisher Scientific) by blotting for 1.5s at 4°C and 100% humidity with a blot force of −5 followed by plunge freezing in liquid ethane. Grids were screened for particle and ice quality at the UBC High Resolution Macromolecular Cryo-Electron Microscopy (HRMEM) facility using a 200kV Glacios TEM (Thermo Fisher Scientific) equipped with a Falcon 3EC DED. All datasets were then collected at the Pacific Northwest Cryo-EM Center (PNCC) using a Titan Krios equipped with a K3 DED and a BioQuantum K3 energy filter with a slit width of 20 eV (Gatan). For the apo p110γ-p101 complex, 6153 super-resolution movies were collected using SerialEM with a total dose of 50e^-^/Å^2^ over 50 frames at a physical pixel size of 1.079Å/pix, using a defocus range of −0.8 to −2μm. For the nanobody-bound p110γ-p101 complex, 6808 super-resolution movies were collected using SerialEM with a total dose of 36.4e^-^ /Å^2^ over 50 frames at a physical pixel size of 1.059Å/pix, using a defocus range of −1 to −2.4μm.

### Cryo-EM image analysis

All data processing was carried out using cryoSPARC v2.18+ unless otherwise specified. For the nanobody-bound p110γ-p101 complex dataset, patch motion correction using default settings was first applied to all movies to align the frames and Fourier crop the outputs by a factor of 2. The contrast transfer function (CTF) of the resulting micrographs was estimated using the patch CTF estimation job with default settings. 2D class averages from a previous dataset were low-pass filtered to 15Å and used as templates to auto-pick 3,762,631 particles, which were then extracted with a box size of 300 pixels. The particles were subjected to 2D classification with the 2D class re-center threshold set to 0.05, and a circular mask of 200Å. 2D class averages that had ice contamination or showed no features were discarded. The remaining 952,705 particles were next used for *ab initio* reconstruction and heterogenous refinement using 2 classes. 692,109 particles from the better 3D reconstruction were curated and any particles from micrographs with a CTF estimation worse than 3Å or total frame motion greater than 30Å were discarded. Per-particle local motion correction was then carried out on the remaining 662,855 particles. The particles were then used for *ab intio* reconstruction and heterogeneous refinement using 4 classes. 320,179 particles from the most complete class were used to carry out homogenous refinement using the 3D reconstruction for that class as a starting model, yielding a reconstruction with an overall resolution of 2.99Å based on the Fourier shell correlation (FSC) 0.143 criterion. The particles were further refined using local CTF refinement before being used for non-uniform refinement with simultaneous global CTF refinement, yielding a map with an overall resolution of 2.90Å. Finally, the map was subjected to a final non-uniform refinement using a mask enveloping the entire volume with the rotation fulcrum centered at the low-resolution nanobody-p101 interaction interface, producing the final map used for model building at a 2.89Å overall resolution.

For the apo p110γ-p101 complex dataset, full-frame motion correction using default settings was first applied to all movies to align the frames. The contrast transfer function (CTF) of the resulting micrographs was estimated using CTFFIND4 with default settings. 2D class averages from a previous dataset were low-pass filtered to 15Å and used as a template to auto-pick 4,792,176 particles, which were then down-sampled by 2 (resulting pixel size of 1.079 Å/pix) and extracted with a box size of 320 pixels. The particles were subjected to multiple rounds of 2D classification with the 2D class re-center threshold set to 0.05, and a circular mask of 200Å. 2D class averages that had ice contamination or did not align to high-resolution were then discarded. The remaining 1,285,510 particles were next subjected to patch CTF estimation and per-particle motion correction before being used for 2 more rounds of 2D classification. 731,169 particles which classified to “good” classes were then used for *ab initio* reconstruction and heterogenous refinement using 2 classes twice iteratively. 196,390 particles from the better 3D reconstruction were used to carry out homogenous refinement using the 3D reconstruction for that class as a starting model, yielding a reconstruction with an overall resolution of 3.49Å based on the Fourier shell correlation (FSC) 0.143 criterion. The map was further refined non-uniform refinement, yielding a map with an overall resolution of 3.36Å.

### Building the structural model of p110γ-p101

The crystal structure of the ΔABD p110γ (PDB: 1E8Y) {Walker:2000bb} was fit into the map using Chimera. Model building was carried out using iterative rounds of automated model building in Phenix, manual model building in COOT (*68*), and refinement in Phenix.real_space_refine using realspace, rigid body, and adp refinement with tight secondary structure restraints (*69*). This allowed for building the ABD, activation loop of the kinase domain, the RBD-C2 and C2-helical linkers in p110γ, and all structured regions of the PBD, and part of the GBD of p101 with high confidence. The regions of the GBD that were manually built included the N-terminal strand (residues 667-676) and the beta hairpin (residues 725 to 745).

Due to the GBD being highly dynamic further automated or manual model building was limited. Starting with this initial model of the GBD, we used a combination of Rosetta de novo modelling (*41*) and folding with trRosetta deep-learned constraints (*42, 43*) to build the remaining GBD (667–837). Rosetta de novo model building was then run on the entire domain starting from this model. This placed an additional strand (residues 677-686). Unfortunately, additional rounds of de novo model-building in Rosetta failed to identify additional regions of the sequence.

Next, we used trRosetta (with some unpublished improvements) to predict contacts for this domain (*42*). The sequence input for trRosetta included the fasta sequence from residues 671-838, and this resulted in ∼2000 aligned sequences, 206 of which (after filtering by 90% of maximum pairwise sequence identity and 50% of minimum coverage) were used to derive constraints. Unfortunately, structure predictions using constraints from trRosetta alone yielded structures that were inconsistent with the observed density data. Therefore, we instead turned to modelling using density and constraints simultaneously. The trRosetta constraints were input along with the density map as inputs to Rosetta comparative modeling (RosettaCM), starting with the partially complete model from Rosetta de novo (*43*). A total of 10,000 modelling trajectories were sampled, yielding good convergence on a model with good agreement to the density (Fig S2). Compared to prior trRosetta modelling procedures (*42*), the constraint weight was reduced to balance the relative contributions of density and the constraints.

To further improve model-map agreement (and overall model geometry), the lowest energy model was used as the input for subsequent rounds of modelling. A total of three rounds of modelling were carried out in this way, each model producing 100 output models, of which the best 10 (by Rosetta + density energies) were carried over to the next round. The final model shows very good agreement with the predicted constraints (Fig S2).

This final model of the GBD was combined with the rest of the p110γ-p101 complex, and final real space refinement was carried out in Phenix using tight secondary structure restraints to give the final model, with full refinement and validation statistics shown in Table S2.

### Expression and Purification of lipidated Gβγ for kinase activity assays

Full length, lipidated human Gβγ (Gβ_1_γ_2_) was expressed in Sf9 insect cells and purified as described previously (*70*). After 65 hours of expression, cells were harvested and the pellets were frozen as described above. Pellets were resuspended in lysis buffer (20 mM HEPES pH 7.7, 100 mM NaCl, 10 mM βME, protease inhibitor (Protease Inhibitor Cocktail Set III, Sigma)) and sonicated for 2 minutes (15s on, 15s off, level 4.0, Misonix sonicator 3000). The lysate was spun at 500 RCF (Eppendorf Centrifuge 5810 R) to remove intact cells and the supernatant was centrifuged again at 25,000 RCF for 1 hour (Beckman Coulter JA-20 rotor). The pellet was resuspended in lysis buffer and sodium cholate was added to a final concentration of 1% and stirred at 4°C for 1 hour. The membrane extract was clarified by spinning at 10,000 RCF for 30 minutes (Beckman Coulter JA-20 rotor). The supernatant was diluted 3 times with NiNTA A buffer (20 mM HEPES pH 7.7, 100 mM NaCl, 10 mM Imidazole, 0.1% C_12_E_10_, 10mM βME) and loaded onto a 5 mL HisTrap™ FF crude column (GE Healthcare) equilibrated in the same buffer. The column was washed with NiNTA A, 6% NiNTA B buffer (20 mM HEPES pH 7.7, 25 mM NaCl, 250 mM imidazole pH 8.0, 0.1% C12E10, 10 mM βME) and the protein was eluted with 100% NiNTA B. The eluent was loaded onto HiTrap^TM^ Q HP anion exchange column equilibrated in Hep A buffer (20 mM Tris pH 8.0, 8 mM CHAPS, 2 mM Dithiothreitol (DTT)). A gradient was started with Hep B buffer (20 mM Tris pH 8.0, 500 mM NaCl, 8 mM CHAPS, 2 mM DTT) and the protein was eluted in ∼50% Hep B buffer. The eluent was concentrated in a 30,000 MWCO Amicon Concentrator (Millipore) to < 1 mL and injected onto a Superdex^TM^ 75 10/300 GL size exclusion column (GE Healthcare) equilibrated in Gel Filtration buffer (20 mM HEPES pH 7.7, 100 mM NaCl, 10 mM CHAPS, 2 mM TCEP). Fractions containing protein were pooled, concentrated, aliquoted, frozen and stored at −80°C.

### Lipid vesicle preparation for kinase activity assays

Lipid vesicles containing 5% brain phosphatidylinositol 4,5-bisphosphate (PI*P*2), 20% brain phosphatidylserine (PS), 50% egg-yolk phosphatidylethanolamine (PE), 10% egg-yolk phosphatidylcholine (PC), 10% cholesterol and 5% egg-yolk sphingomyelin (SM) were prepared by mixing the lipids solutions in organic solvent. The solvent was evaporated in a stream of argon following which the lipid film was desiccated in a vacuum for 45 minutes. The lipids were resuspended in lipid buffer (20 mM HEPES pH 7.0, 100 mM NaCl and 10 % glycerol) and the solution was sonicated for 15 minutes. The vesicles were subjected to five freeze thaw cycles and extruded 11 times through a 100-nm filter (T&T Scientific: TT-002-0010). The extruded vesicles were sonicated again for 5 minutes, aliquoted and stored at −80°C. Final vesicle concentration was 5 mg/mL.

### Lipid kinase activity assays

All lipid kinase activity assays employed the Transcreener ADP2 Fluorescence Intensity (FI) Assay (Bellbrook labs) which measures ADP production. For assays comparing the activities of p110γ, p110γ-p101 and p110γ-p84, PM-mimic vesicles at a final concentration of 1 mg/mL, ATP at a final concentration of 100 μM ATP and Gβγ at 1.5 μM final concentration were used. Final concentration of kinase was 3000nM for all basal conditions. For conditions with Gβγ, p110γ: 3000 nM, p110γ-p84: 1000 nM and p110γ-p101: 30 nM. 2 μL kinase solution at 2X final concentration was mixed with 2 μL substrate solution containing ATP, vesicles and Gβγ or Gβγ gel filtration buffer and the reaction was allowed to proceed for 60 minutes at 20°C. For assays comparing mutants, kinase was mixed with vesicles at 1mg/mL, ATP at 100 μM and Gβγ at 75 nM or 1.5 μM final concentrations and the reaction was allowed to proceed for 60 minutes at 37°C. The reactions were stopped with 4 μL of 2X stop and detect solution containing Stop and Detect buffer, 8 nM ADP Alexa Fluor 594 Tracer and 93.7 μg/mL ADP2 Antibody IRDye QC-1 and incubated for 50 minutes. The fluorescence intensity was measured using a SpectraMax M5 plate reader at excitation 590 nm and emission 620 nm. This data was normalized against the measurements obtained for 100 μM ATP and 100 μM ADP. The % ATP turnover was interpolated from a standard curve (0.1-100 μM ADP). This was then used to calculate the specific activity of the enzyme.

For assays measuring nanobody inhibition at saturating Gβγ concentrations, kinase at 4X concentration was mixed with nanobody at 4X concentration to obtain 2X enzyme-nanobody solution (6 μM final nanobody). Final concentration of kinase was 3000nM for all basal conditions. For conditions with Gβγ, p110γ-p84: 1500 nM and p110γ-p101: 50 nM. 2 μL of this solution was mixed with 2 μL of 2X substrate solution containing ATP (100 μM final), vesicles (1 mg/mL final) with or without Gβγ (1.5 μM final) to start the reaction and allowed to proceed for 60 minutes at 20 °C. Following this, the reaction was stopped, the intensity was measured, the data was normalized and specific activity calculated as described above. For the nanobody IC_50_ curve, kinase at 4X concentration was mixed with nanobody at 4X concentration to obtain 2X enzyme-nanobody solution (200nM final kinase; 41-10,000 nM final nanobody). 2 μL of this solution was mixed with 2 μL of 2X substrate solution containing ATP (100 μM final), vesicles (2 mg/mL final) and Gβγ (600 nM final) to start the reaction and allowed to proceed for 60 minutes at 20 °C. Following this, the reaction was stopped, the intensity was measured, and the data was normalized as described above. The normalized values for conditions with nanobody were then further normalized against the condition with maximal activity (no nanobody).

### Bio-layer interferometry

Biolayer interferometry was performed using Octet K2 (Fortebio, Inc.). His-tagged nanobody was immobilized on an Anti-Penta-His biosensor for 600 seconds and the sensor was dipped into solutions of p110γ-p101 at 1.5, 6 and 25 nM final concentrations diluted in kinetics buffer (KB) containing 20mM Tris pH 8.5, 100mM NaCl, 50mM ammonium sulphate, 0.1% bovine serum albumin and 0.02% tween-20. The association step was allowed to proceed for 600 seconds followed by a dissociation step in KB without protein for 600 seconds. The baseline was obtained by dipping sensor without nanobody into a solution containing 25 nM p110γ-p101 in a similar fashion. The average K_d_ was calculated from the three binding curves based on their global fit to a 1:1 binding model. *Hydrogen Deuterium Exchange Mass Spectrometry (HDX-MS) (STAR methods)*

## HDX-MS sample preparation

For HDX reactions comparing p110γ alone and p110γ in complex with p101 or p84, exchange was carried out in a 50 µl reaction containing 20 picomoles of protein, either p110γ, p110γ-p84 or p110γ-p101. To initiate hydrogen-deuterium exchange, 1.5µL of either protein was incubated with 48.5 µL of D_2_O buffer solution (20mM HEPES pH 7.5, 100mM NaCl, 94.3% D_2_O) for five time points (3s on ice, 3s, 30s, 300s, 3000s at room temperature) to give a final concentration of 91.5% D_2_O.

HDX reactions comparing full length p110γ and ABD truncated p110γ were conducted in 50 µl reaction volumes with a final p110γ amount of 15 pmol. Exchange was carried out for four time points (3s, 30s, 300s and 3000s at room temperature). To initiate hydrogen-deuterium exchange, 1.2 µL of either protein was incubated with 48.8 µL of D_2_O buffer solution (20mM HEPES pH 7.5, 100mM NaCl, 94.3% D_2_O) to give a final concentration of 92% D_2_O.

HDX reactions comparing p110γ-p101 with and without nanobody were conducted in 50 µl reaction volumes with a final p110γ amount of 16 pmol and a final nanobody amount of 7 pmol. Exchange was carried out for two time points (3s and 300s at room temperature). To initiate hydrogen-deuterium exchange, 1µL of p110γ-p101 and 1µL of nanobody was incubated with 48 µL of D_2_O buffer solution (20mM HEPES pH 7.5, 100mM NaCl, 94.3% D_2_O) to give a final concentration of 90.5% D_2_O.

HDX reactions comparing wild-type p110γ-p101, R472C p110γ-p101, and E347K p110γ-p101 were conducted in 50 µl reaction volumes with a final protein amount of 12.5 pmol. Exchange was carried out for five time points (3s on ice, 3s, 30s, 300s and 3000s at room temperature). To initiate hydrogen-deuterium exchange, 2 µL of protein was incubated with 48 µL of D_2_O buffer solution (20mM HEPES pH 7.5, 100mM NaCl, 94.3% D_2_O) to give a final concentration of 90.5% D_2_O. All exchange reactions were terminated by the addition of ice-cold quench buffer to give a final concentration 0.6M guanidine-HCl and 0.9% formic acid and samples were frozen in liquid nitrogen and stored at −80°C. All experiments were carried out in independent triplicate.

## Protein digestion and MS/MS data collection

Protein samples were rapidly thawed and injected onto an integrated fluidics system containing a HDx-3 PAL liquid handling robot and climate-controlled chromatography system (LEAP Technologies), a Dionex Ultimate 3000 UHPLC system, as well as an Impact HD QTOF Mass spectrometer (Bruker). The full details of the fluidics system are described in (*71*). The protein was run over either one (at 10°C) or two (at 10°C and 2°C) immobilized pepsin columns (Applied Biosystems; Poroszyme Immobilized Pepsin Cartridge, 2.1 mm x 30 mm; Thermo-Fisher 2-3131-00; Trajan; ProDx protease column, 2.1 mm x 30 mm PDX.PP01-F32) at 200 µL/min for 3 minutes. The resulting peptides were collected and desalted on a C18 trap column (Acquity UPLC BEH C18 1.7mm column (2.1 x 5 mm); Waters 186003975). The trap was subsequently eluted in line with a C18 reverse-phase separation column (Acquity 1.7 mm particle, 100 x 1 mm^2^ C18 UPLC column, Waters 186002352), using a gradient of 5-36% B (Buffer A 0.1% formic acid; Buffer B 100% acetonitrile) over 16 minutes. Mass spectrometry experiments acquired over a mass range from 150 to 2200 m/z using an electrospray ionization source operated at a temperature of 200°C and a spray voltage of 4.5 kV.

## Peptide identification

Peptides were identified from the non-deuterated samples of p110γ alone, p110γ-p84, or p110γ-p101 using data-dependent acquisition following tandem MS/MS experiments (0.5 s precursor scan from 150-2000 m/z; twelve 0.25 s fragment scans from 150-2000 m/z). MS/MS datasets were analyzed using PEAKS7 (PEAKS), and peptide identification was carried out by using a false discovery-based approach, with a threshold set to 1% using a database of purified proteins and known contaminants (*72*). The search parameters were set with a precursor tolerance of 20 ppm, fragment mass error 0.02 Da, charge states from 1-8, with a selection criterion of peptides that had a −10logP score of 21.7.

## Mass Analysis of Peptide Centroids and Measurement of Deuterium Incorporation

HD-Examiner Software (Sierra Analytics) was used to automatically calculate the level of deuterium incorporation into each peptide. All peptides were manually inspected for correct charge state, correct retention time, appropriate selection of isotopic distribution, etc. Deuteration levels were calculated using the centroid of the experimental isotope clusters. The results for these proteins are presented as the raw percent deuterium incorporation, as shown in Supplemental Data, with the only correction being applied correcting for the deuterium oxide percentage of the buffer utilized in the exchange (91.7% for comparing p110γ to p110γ-p101 and p110γ-p84, 86.8% for comparing p110γ-p101 with NB1-PIK3R5, 92% for the ABD deletion, and 90.5% for the oncogenic mutants). No corrections for back exchange that occurs during the quench and digest/separation were applied. Attempts to generate a fully deuterated class I PI3K sample were unsuccessful, which is common for large macromolecular complexes. Therefore, all deuterium exchange values are relative.

Differences in exchange in a peptide were considered significant if they met all three of the following criteria: ≥5% change in exchange, ≥0.4 Da difference in exchange, and a p value <0.01 using a two tailed student t-test. The raw HDX data are shown in two different formats.

The raw peptide deuterium incorporation graphs for a selection of peptides with significant differences are shown, with the raw data for all analyzed peptides in the source data. To allow for visualization of differences across all peptides, we utilized number of deuteron difference (#D) plots. These plots show the total difference in deuterium incorporation over the entire H/D exchange time course, with each point indicating a single peptide. These graphs are calculated by summing the differences at every time point for each peptide and propagating the error (example Fig S6A-B). For a selection of peptides we are showing the %D incorporation over a time course, which allows for comparison of multiple conditions at the same time for a given region (Fig. S6C). Samples were only compared within a single experiment and were never compared to experiments completed at a different time with a different final D_2_O level. The data analysis statistics for all HDX-MS experiments are in Table S3 according to the guidelines of (*73*). The mass spectrometry proteomics data have been deposited to the ProteomeXchange Consortium via the PRIDE partner repository (*74*) with the dataset identifier PXD025209.

### Total Internal Reflection Fluorescence Microscopy (TIRF) (STAR methods)

#### Purification of recombinant farnesyl G_β1_Gγ_2_ and SNAP-G_β1_Gγ_2_ for TIRF microscopy

Genes encoding bovine G_β1_ and Gγ_2_ were cloned into baculovirus expression vectors Gibson assembly. This was achieved using YFP-G_β1_ (Addgene plasmid # 36397) and YFP-Gγ_2_ (Addgene plasmid # 36102) as templates for PCR. These G_β1_ and Gγ_2_ containing plasmids were kindly provided to Addgene by Narasimhan Gautam (*75*). We recombinantly expressed either G_β1_/his_6_-TEV-Gγ_2_ or SNAP-G_β1_/his_6_-TEV-Gγ_2_ in High five insect cells using a dual expression vector system with tandem polyhedron promoters. General procedures for making BACMID and baculovirus were performed as previously described (*76*). For G_β1_/his_6_-TEV-Gγ_2_ or SNAP-G_β1_/his_6_-TEV-Gγ_2_ expression, 2-4 liters of High Five cells (2 x 10^6^ cells/mL) were infected with 2% vol/vol of baculovirus. Cultures were then grown in 3 liter shaker flasks (120 rpm) for 48 hours at 27 °C before harvesting. Insect cells were harvested by centrifugation and stored as 10 g pellets in the −80°C until initiating the purification.

To isolate farnesylated G_β1_/his_6_-TEV-Gγ_2_ or SNAP-G_β1_/his_6_-TEV-Gγ_2_ complexes from insect cells, we used a hybrid purification protocol based on both (*70, 77*). Cells were lysed by dounce homogenization into the buffer containing 50 mM HEPES-NaOH [pH 8], 100 mM NaCl, 3 mM MgCl_2_, 0.1 mM EDTA, 10 µM GDP, 10 mM BME, Sigma PI tablets, 1 mM PMSF, DNase. Cell lysate was subjected to low speed centrifugation (10 min at 800 RCF) to remove nuclei and large cell debri. Next, soluble cell membranes isolated from partially clarified lysate were pelleted by centrifugation using a Beckman Ti45 rotor (100,000 RCF for 30 minutes). Pellets containing the membrane fraction of the cell lysate were solubilize in membrane extraction buffer: 50 mM HEPES-NaOH (pH 8), 50 mM NaCl, 3 mM MgCl_2_, 1% sodium deoxycholate (wt/vol, Sigma D6750), 10µM GDP (Sigma G7127), 10 mM BME, and Sigma protease inhibitor tablet in order to solubilize farnesylated G_β1_/his_6_-TEV-Gγ_2_ or SNAP-G_β1_/his_6_-TEV-Gγ_2_. Membranes were resuspended in enough membrane extraction buffer to reach a protein concentration of 5 mg/mL. We used a dounce homogenizer to break apart the membrane followed by stirring in a beaker at 4°C for 1 hour. Following membrane extraction, insoluble material was removed by 4°C ultracentrifugation at 100,000 RCF for 45 minutes. Clarify supernatant containing detergent solubilized G_β1_/his_6_-TEV-Gγ_2_ or SNAP-G_β1_/his_6_-TEV-Gγ_2_ was diluted 5-fold in post membrane extraction buffer: 20 mM HEPES-NaOH (pH 7.7), 100 mM NaCl, 0.1 % C_12_E_10_ (Polyoxyethylene (10) lauryl ether; Sigma, P9769), 25 mM imidazole, and 2 mM BME. For affinity purification of G_β1_/his_6_-TEV-Gγ_2_ or SNAP-G_β1_/his_6_-TEV-Gγ_2_, 7-10 mL of Qiagen NiNTA resin (50% slurry) was added diluted post membrane extraction sample and incubated on a stir plate in a beaker at 4°C for 2 hours. Using a gravity flow column, NiNTA resin was washed with 20 column volumes of buffer containing 20 mM HEPES-NaOH (pH 7.7), 100 mM NaCl, 0.1 % C_12_E_10_, 20 mM imidazole, 2 mM BME. Next, the contaminating G alpha subunit of the heterotrimeric G-protein complex was eluted by washing with warm buffer (30°C) containing 20 mM HEPES-NaOH (pH 7.7), 100 mM NaCl, 0.1 % C_12_E_10_, 20 mM imidazole, 2 mM BME, 50 mM MgCl_2_, 10µM GDP, 30 µM AlCl_3_, 10 mM NaF. When dissolving AlCl_3_ into water we worked in a fume hood to prevent inhalation of gaseous HCl. Finally, G_β1_/his_6_-TEV-Gγ_2_ or SNAP-G_β1_/his_6_-TEV-Gγ_2_ were eluted from NiNTA resin with buffer containing 20 mM Tris-HCl (pH 8.0), 25 mM NaCl, 0.1 % C_12_E_10_, 200 mM imidazole, 2 mM BME. Eluate was combined with TEV protease and incubated overnight in the NTA elution buffer at 4°C. During day two of the purification, G_β1_Gγ_2_ or SNAP-G_β1_Gγ_2_ were desalted into buffer containing 20 mM Tris-HCl (pH 8.0), 25mM NaCl, 8 mM CHAPS, 2 mM TCEP and loaded onto a MonoQ column equilibrated with the same buffer. G_β1_Gγ_2_ or SNAP-G_β1_Gγ_2_ eluted from the MonoQ column in the presence of 175-200 mM NaCl. A single peak was combined and concentrated using a Millipore Amicon Ultra-4 (10 kDa MWCO) centrifuge filter. G_β1_Gγ_2_ or SNAP-G_β1_Gγ_2_ were respectively loaded on either Superdex 75 or Superdex 200 gel filtration columns equilibrated 20 mM Tris [pH 8.0], 100 mM NaCl, 8 mM CHAPS, 2 mM TCEP. We pooled peak fractions and concentrated the protein using a Millipore Amicon Ultra-4 (10 kDa MWCO) centrifuge tube. Concentrated G_β1_Gγ_2_ or SNAP-G_β1_Gγ_2_ was aliquoted and flash frozen liquid nitrogen. Using single molecule TIRF microscopy, we determined that the quality of the protein was similarly high when the protein was frozen in the absence or presence of 10% glycerol.

#### Fluorescent labeling of ybbr-p110γ and p110γ/ybbr-p101

The Dyomics647-CoA derivative was generated in-house by combining 15 mM Dyomics647 maleimide (Dyomics, Cat #647P1-03) in DMSO with 10 mM CoA (Sigma, #C3019, MW = 785.33 g/mole) dissolved in 1x PBS. This mixture was incubated overnight at 23°C. Unreacted Dyomics647 maleimide was quenched by the addition of 5 mM DTT. DY647-CoA can be stored at −20°C for at least one year. We labeled recombinant p110γ/p101 containing a N-terminal ybbR13 motif (D<u>S</u>LEFIASKLA) using Sfp transferase and DY647-CoA (*78*). Chemical labeling was achieved by combining 5 µM p110γ/ybbr-p101 (or ybbr-p110γ), 4 µM Sfp-his_6_, and 10 µM DY647-CoA, in 2 mL of buffer containing 20 mM Tris [pH 8], 150 mM NaCl, 10 mM MgCl_2_, 10% Glycerol, 1 mM TCEP, 0.05% CHAPS. Following a 4 hour labeling reaction on ice, excess DY647-CoA was removed using a gravity flow PD-10 column. p110γ/Dy647-ybbr-p101 (or Dy647-ybbr-p110γ) was concentrated in a 50 kDa MWCO Amicon centrifuge tube and loaded on a Superdex 200 gel filtration column equilibrated in 20 mM Tris [pH 8], 150 mM NaCl, 10% glycerol, 1 mM TCEP, 0.05% CHAPS. Peak fractions were pooled and concentrated to 5-10µM before flash freezing with liquid nitrogen.

#### Preparation of supported lipid bilayers

The following lipids were used to generated small unilamellar vesicles (SUVs) and subsequently supported lipid bilayers: 1,2-dioleoyl-sn-glycero-3-phosphocholine (18:1 DOPC, Avanti # 850375C) and 1,2-dioleoyl-*sn-*glycero-3-phospho-L-serine (18:1 DOPS, Avanti # 840035C). To make liposomes, 2 µmoles total lipids are combined in a 35 mL glass round bottom flask containing 2 mL of chloroform. Lipids are dried to a thin film using rotary evaporation with the glass round-bottom flask submerged in a 42°C water bath. After evaporating all the chloroform, the round bottom flask was flushed with nitrogen gas for at least 30 minutes. Resuspend lipid film in 2 mL of PBS [pH 7.2], making a final concentration of 1 mM total lipids. All lipid mixtures expressed as percentages (e.g. 95% DOPC, 5% DOPS) are equivalent to molar fractions. To generate 30-50 nm SUVs, 1 mM total lipid mixtures were extruded through a 0.03 µm pore size 19 mm polycarbonate membrane (Avanti #610002) with filter supports (Avanti #610014) on both sides of the PC membrane.

Supported lipid bilayers are formed on 25×75 mm coverglass (IBIDI, #10812). Coverglass was first cleaned with 2% Hellmanex III (Fisher, Cat#14-385-864) heated to 60-70^0^C in a glass coplin jar. Incubate for at least 30 minutes. Wash coverglass extensively with MilliQ water and then etched with Pirahna solution (1:3, hydrogen peroxide:sulfuric acid) for 10-15 minutes the same day SLBs were formed. Etched coverglass, in water, is rapidly dried with nitrogen gas before adhering to a 6-well sticky-side chamber (IBIDI, Cat# 80608). Form SLBs by flowing 30 nm SUVs diluted in PBS [pH 7.2] to a total lipid concentration of 0.25 mM. After 30 minutes, IBIDI chambers are washed with 5 mL of PBS [pH 7.2] to remove non-absorbed SUVs. Membrane defects are blocked for 15 minutes with a 1 mg/mL beta casein (Thermo FisherSci, Cat# 37528) diluted in 1x PBS [pH 7.4]. Before use as a blocking protein, frozen 10 mg/mL beta casein stocks were thawed, centrifuged for 30 minutes at 21370 x *g*, and 0.22 µm syringe filtered. After blocking SLBs with beta casein, membranes were washed again with 1mL of PBS, followed by 1 mL of TIRF-M imaging buffer before TIRF-M.

#### Single molecule TIRF microscopy

All supported membrane TIRF-M experiments were performed using the following reaction buffer: 20 mM HEPES [pH 7.0], 150 mM NaCl, 1 mM ATP, 5 mM MgCl_2_, 0.5 mM EGTA, 20 mM glucose, 200 µg/mL beta casein (ThermoScientific, Cat# 37528), 20 mM BME, 320 µg/mL glucose oxidase (Serva, #22780.01 *Aspergillus niger*), 50 µg/mL catalase (Sigma, #C40-100MG Bovine Liver), and 2 mM Trolox (*76*). Perishable reagents (i.e. glucose oxidase, catalase, and Trolox) were added 5-10 minutes before image acquisition.

Single-molecule imaging experiments were performed on an inverted Nikon Ti2 microscope using a 100x Nikon objective (1.49 NA) oil immersion TIRF objective. We manually control the x-axis and y-axis positions using an Nikon motorized stage and joystick. Fluorescently labelled proteins were excited with either a 488 nm or 637 nm diode laser (OBIS laser diode, Coherent Inc. Santa Clara, CA) controlled with a Vortran laser drive with acousto-optic tuneable filters (AOTF) control. The power output measured through the objective for single particle imaging was 1-3 mW. Excitation light was passed through the following dichroic filter cubes before illuminating the sample: (1) ZT488/647rpc and (2) ZT561rdc (ET575LP) (Semrock). Fluorescence emission was detected on an iXion Life 897 EMCCD camera (Andor Technology Ltd., UK) after passing through a Nikon Ti2 emission filter wheel containing the following 25 mm emission filters: ET525/50M, ET600/50M, ET700/75M (Semrock). All experiments were performed at room temperature (21-23^0^C). Microscope hardware was controlled with Nikon NIS elements.

#### Single particle tracking

After the 16-bit .tif images were cropped down to 400×400 to minimize differences in field illumination. Upon starting the TrackMate plugin (*79*) on ImageJ/Fiji, the initial calibration settings were default and enter ‘YES’ for allow swapping of Z or T (depth or time). Select the LoG detector and enter the appropriate protein-related dimensions. Check the boxes applying median filter and sub-pixel localization. Apply default settings for initial thresholding, run hyperstack display, and then filter particles based on mean intensity. Choose a tracker (simple LAP tracker was used in this paper, but LAP tracker also works). Set the following filters on the tracks: Track Start (remove particles at start of movie); Track End (remove particles at end of movie); Track displacement; X-Y location (above and below totaling 4 filters). Continue running program and display options according to one’s preference. From the generated files, the dwell times were extracted using a custom MATLAB script. A histogram and cumulative frequency distributions were generated for each data set. To calculate the dwell times for membrane bound lipid kinases we sorted into a cumulative distribution frequency (CDF) plot with the frame interval as the bin (e.g. 28 ms). A typical histogram contains dwell time from n = 2000-3000 tracked particles from n = 3 movies. The log_10_(1-CDF) is then plotted against the dwell time and fit to a single or double exponential. For double exponential fit, alpha represents the percentage of the fast dissociating molecules characterized by τ_1_.

### Single exponential model

*f*(*t*) = *e*^(-*x*/⁄τ)^

### Two exponential model

*f*(*t*) = *a* ∗ *e*^(-x/⁄τ)^+(1 − *a*) ∗ *e*^(-x/⁄τ2)^

## Data availability statement

The electron microscopy data has been deposited in the electron microscopy data bank with accession numbers (EMDB: 23808 [p110γ-p101-NB1-PIK3R5] and 23812 [p110γ-p101]) and associated structural models have been deposited to the protein data bank with accession numbers (PDB: 7MEZ). The mass spectrometry proteomics data have been deposited to the ProteomeXchange Consortium via the PRIDE (*74*) partner repository with the dataset identifier PXD025209. All raw data in all figures are available in the source data.

## Acknowledgements

J.E.B. is supported the Canadian Institute of Health Research (CIHR, 168998), a Michael Smith Foundation for Health Research (MSFHR, scholar 17686), and the Cancer Research Society (CRS-24368). C.K.Y. is supported by CIHR (FDN-143228) and the Natural Sciences and Engineering Research Council of Canada (RGPIN-2018-03951). S.D.H. is supported by a NSF CAREER Award (MCB-2048060). F.D. is supported by NIH-GM123089. JS and EP acknowledge the support and the use of resources of Instruct-ERIC, part of the European Strategy Forum on Research Infrastructures (ESFRI), and the Research Foundation - Flanders (FWO). This research project was supported in part by the UBC High Resolution Macromolecular Cryo-Electron Microscopy Facility (HRMEM). A portion of this research was supported by NIH grant U24GM129547 and performed at the PNCC at OHSU and accessed through EMSL (grid.436923.9), a DOE Office of Science User Facility sponsored by the Office of Biological and Environmental Research. We appreciate help from Theo Humphreys and Rose Marie Haynes with data collection at PNCC.

## Supplemental Figures and Tables

**Fig S1.**
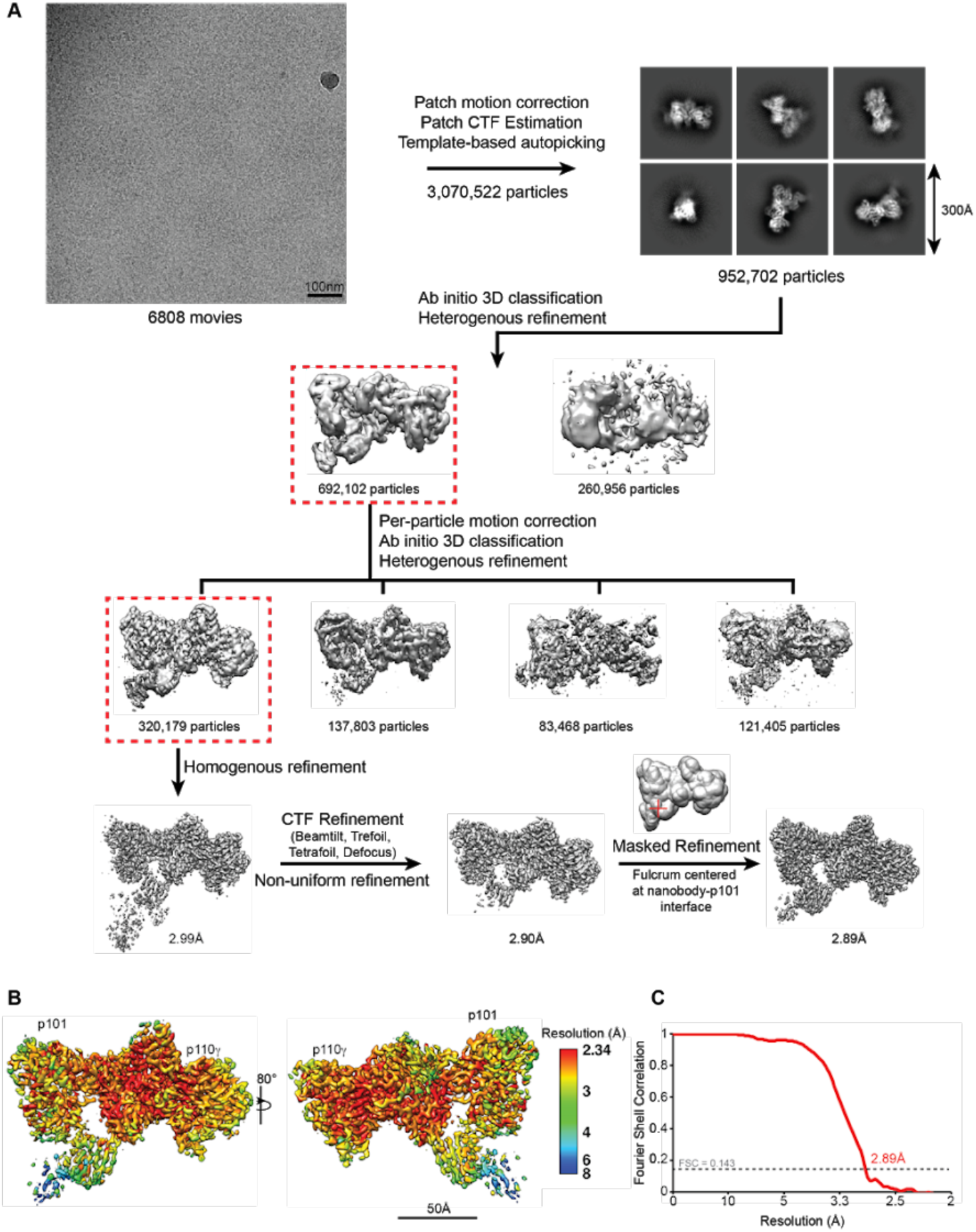
Cryo-EM analysis for the p110γ-p101-nanobody complex. **A.** Cryo-EM analysis workflow showing a representative micrograph, representative 2D class averages and image processing strategy used for the 3D reconstruction of the p110γ-p101 nanobody complex. **B.** p110γ-p101 reconstruction coloured according to local resolution as estimated using cryoSPARC v3.1. **C.** Gold-standard fourier shell correlation (FSC) curve after auto-tightening by cryoSPARC for the final p110γ-p101 map.

**Fig. S2.**
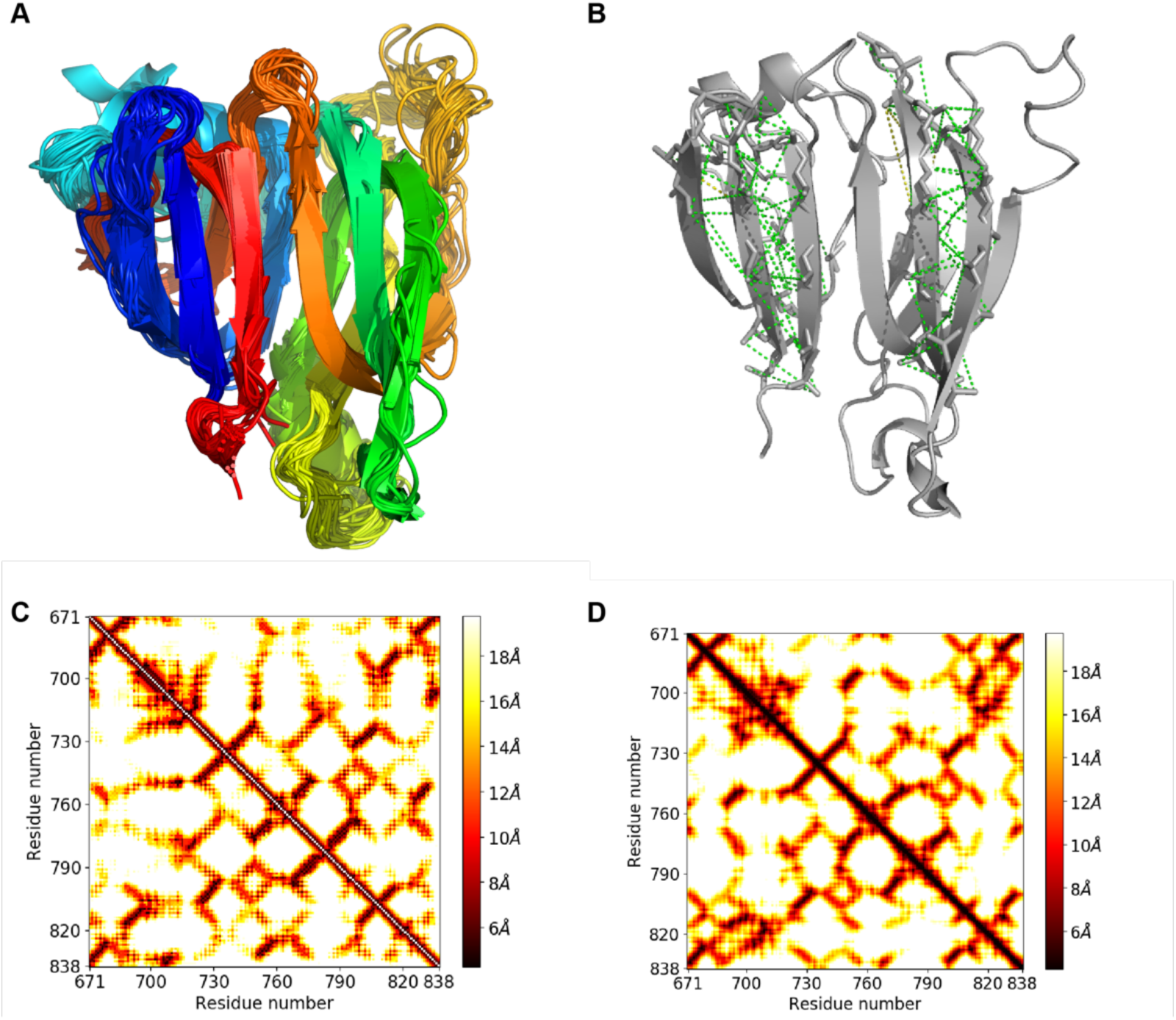
trRosetta modelling of the GBD of p101. **A.** An overlay of the ensemble of 100 models from our final model building in RosettaCM. A majority of positions show tightly overlapping chain trace, with all models converging on the same overall topology. **B.** Residues predicted by trRosetta with >99% probability to have CB atoms within 10 angstroms distance, mapped onto the final model. Green indicates that the actual distance is within 10 angstroms, yellow within 12 angstroms, and red for >12 angstroms. **C. and D.** The predicted and actual distances between beta-carbons of residues in the p101 nanobody-interacting region. The colour of each pixel corresponds to the distance in angstroms between these atoms. Plotted on the left is the least distance predicted by our improved trRosetta pipeline with >95% probability for each pair of CB atoms. On the right are the actual distances between these atoms. Our improved trRosetta pipeline correctly predicts beta-strand interactions between residues 720 and 740, 750 and 770, as well as 770 and 800, among others.

**Figure S3.**
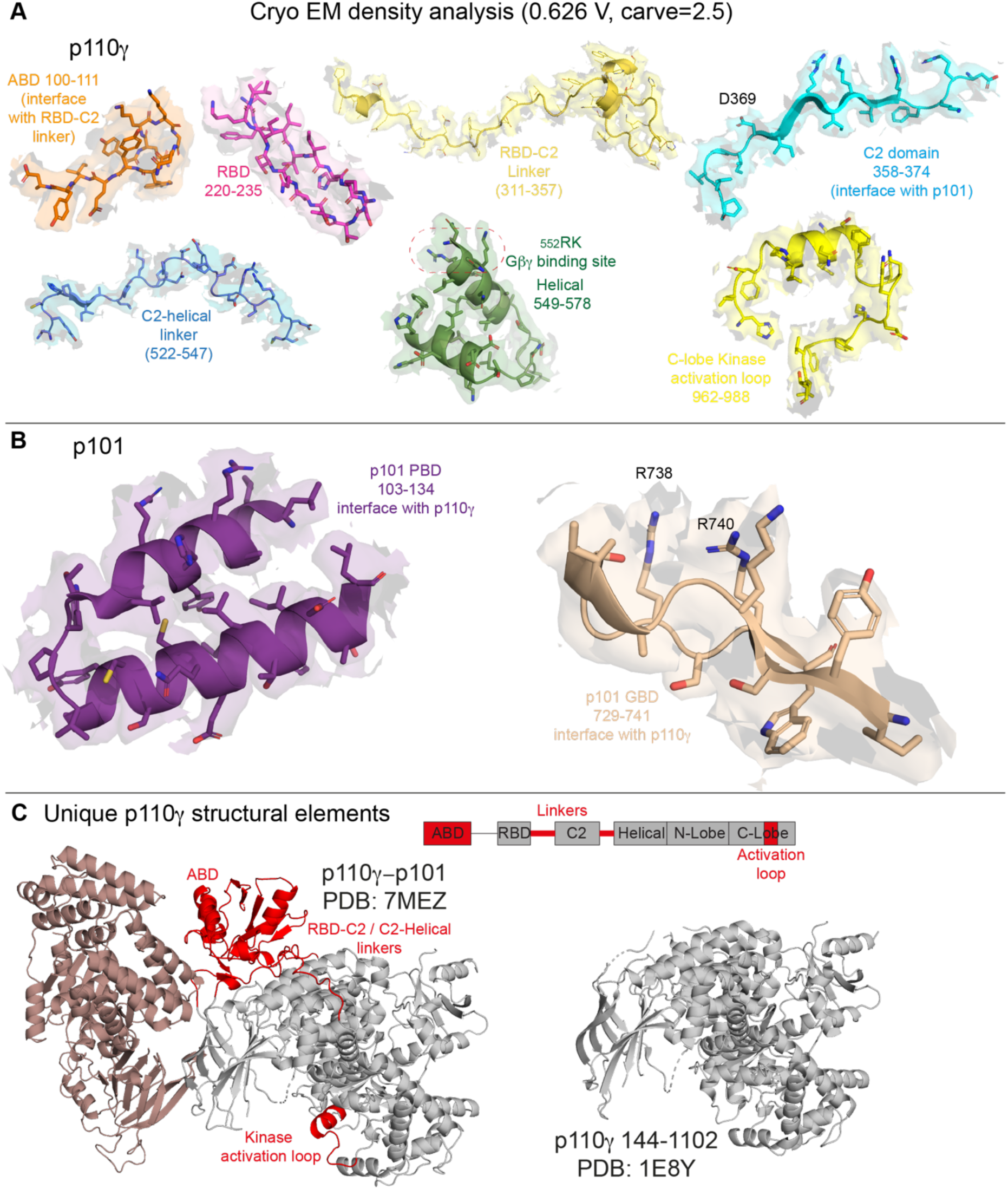
Structural analysis of the p110γ-p101 model. **A.** Electron density in select regions covering all five domains of p110γ **B.** Electron density in select regions of the PBD and GBD of p101 **C.** Novel structural features in the p110 subunit (in red) that were previously absent in the structure of p110γ (144-1102).

**Fig S4.**
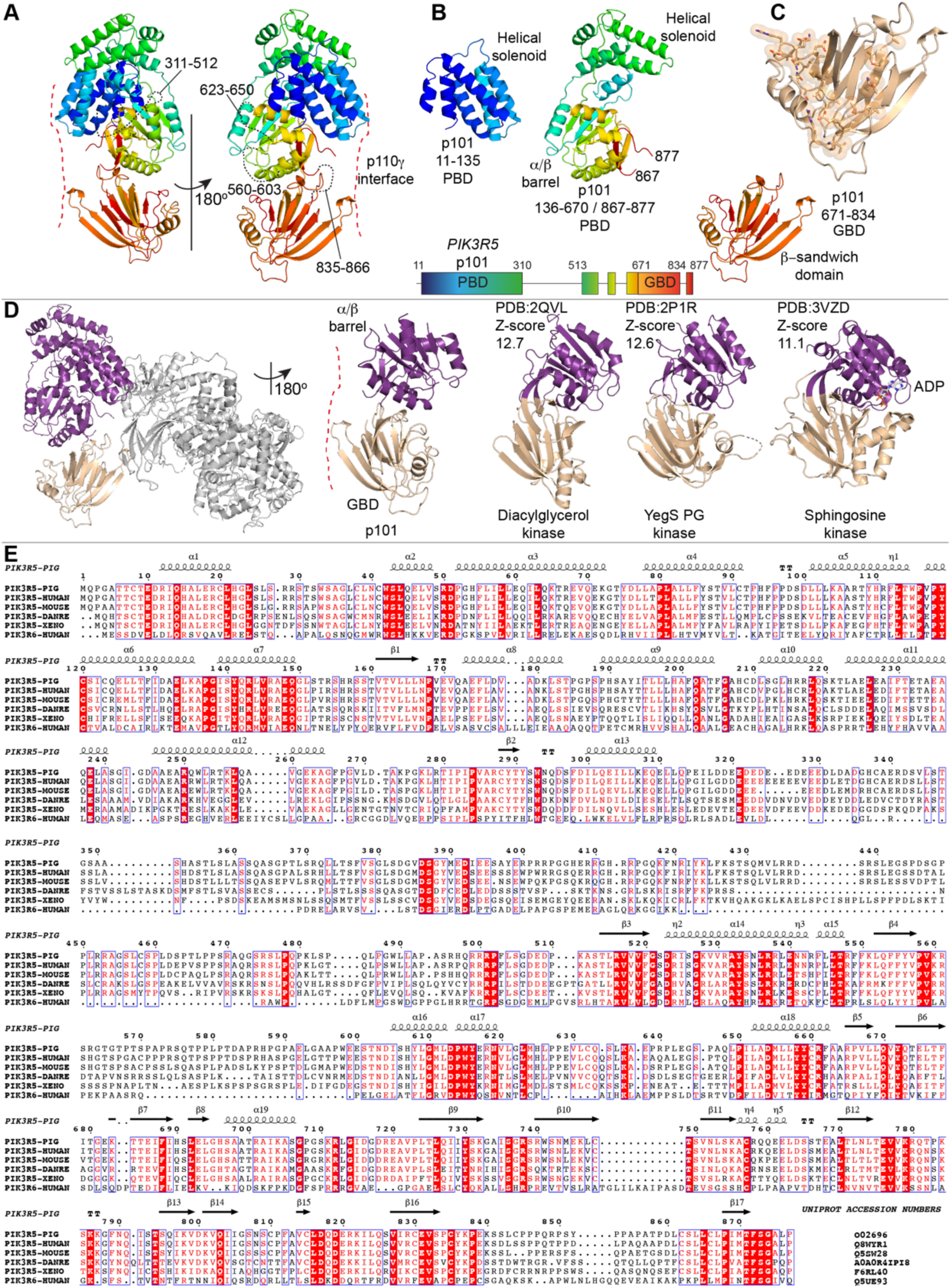
p101 (*PIK3R5*) protein: Structure, structural conservation with DGKs, evolutionary conservation of p101, and comparison with p84 (*PIK3R6*) **A.** Domain map and model of p101 coloured from N-to C-terminus in the rainbow spectrum from blue through red. **B.** Representation of different regions of the model in panel A showing various protein folds in the PBD and GBD domains of p101. **C.** Zoom in on the GBD, showing residues identified as important in Gβγ activation of p110γ-p101 as sticks/spheres. **D.** Structural comparison of the α/β barrel and β-sandwich in p101 compared to the corresponding regions in diacylglycerol kinase, sphingosine kinase 1, and the *Salmonella* phosphatidylglycerol kinase YegS. **E.** Alignment (generated with ESPript 3.0) showing evolutionary conservation of residues in porcine p101 with p101 sequences from human, mouse, Xenopus and zebrafish and p84 sequence from human. The secondary structure elements of porcine p101 are shown above the alignment.

**Fig S5.**
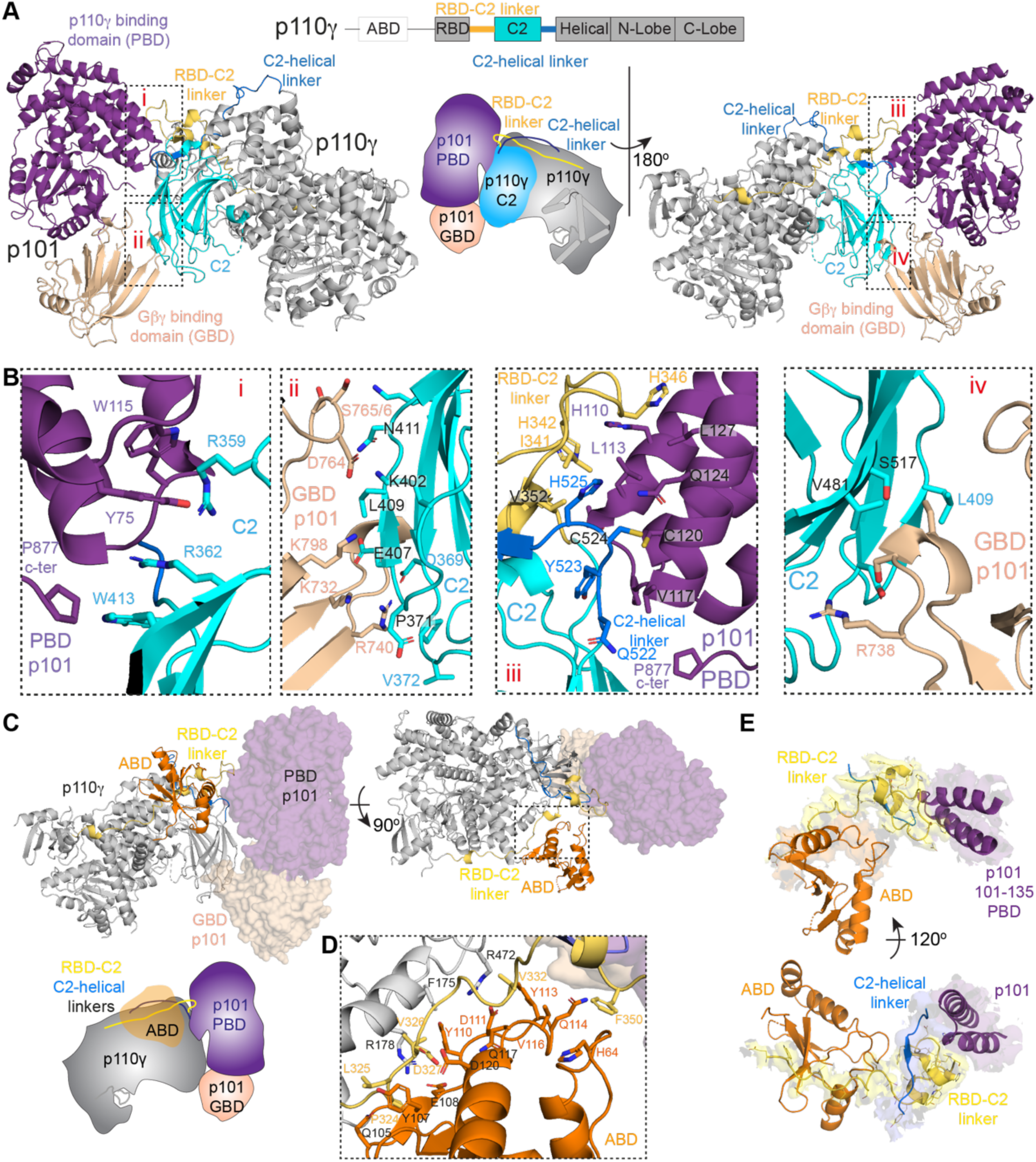
Interface details for p110γ with p101 and ABD. **A.** Cartoon representation of the p110γ-p101 complex, with p101 colored as in Figure 1, and p110γ colored according to the attached schematic, with p101 interacting regions (RBD-C2 linker, C2, and the C2-helical linker) indicated. Important features are shown in a cartoon schematic. Interacting regions are indicated in the boxes, and are labeled i-iv. **B.** Residues that mediate the interaction between p110γ and p101. Residues that have more than 20 Å of buried surface area are labelled and shown in a stick representation. **C.** The structure of p110γ-p101 complex, with p110γ shown as cartoon, and p101 as a transparent surface. The different domains are colored as indicated according to the cartoon schematic. **D.** Residues that mediate the interaction between the p110γ ABD and the rest of p110γ. Residues that have more than 20 Å of buried surface area are labelled and shown in a stick representation. **E.** The ABD of p110γ coordinates the RBD-C2 linker of p110γ to interact with p101. The RBD-C2 linker, ABD, and the region of p101 that binds the RBD-C2 linker are shown in a cartoon representation. The electron density of the RBD-C2 linker and region of p101 that binds the RBD-C2 linker are visible (both views), with the ABD interface with the RBD-C2 linker (top view) and the N-terminus of the C2-helical linker domain (bottom view) are shown.

**Fig S6.**
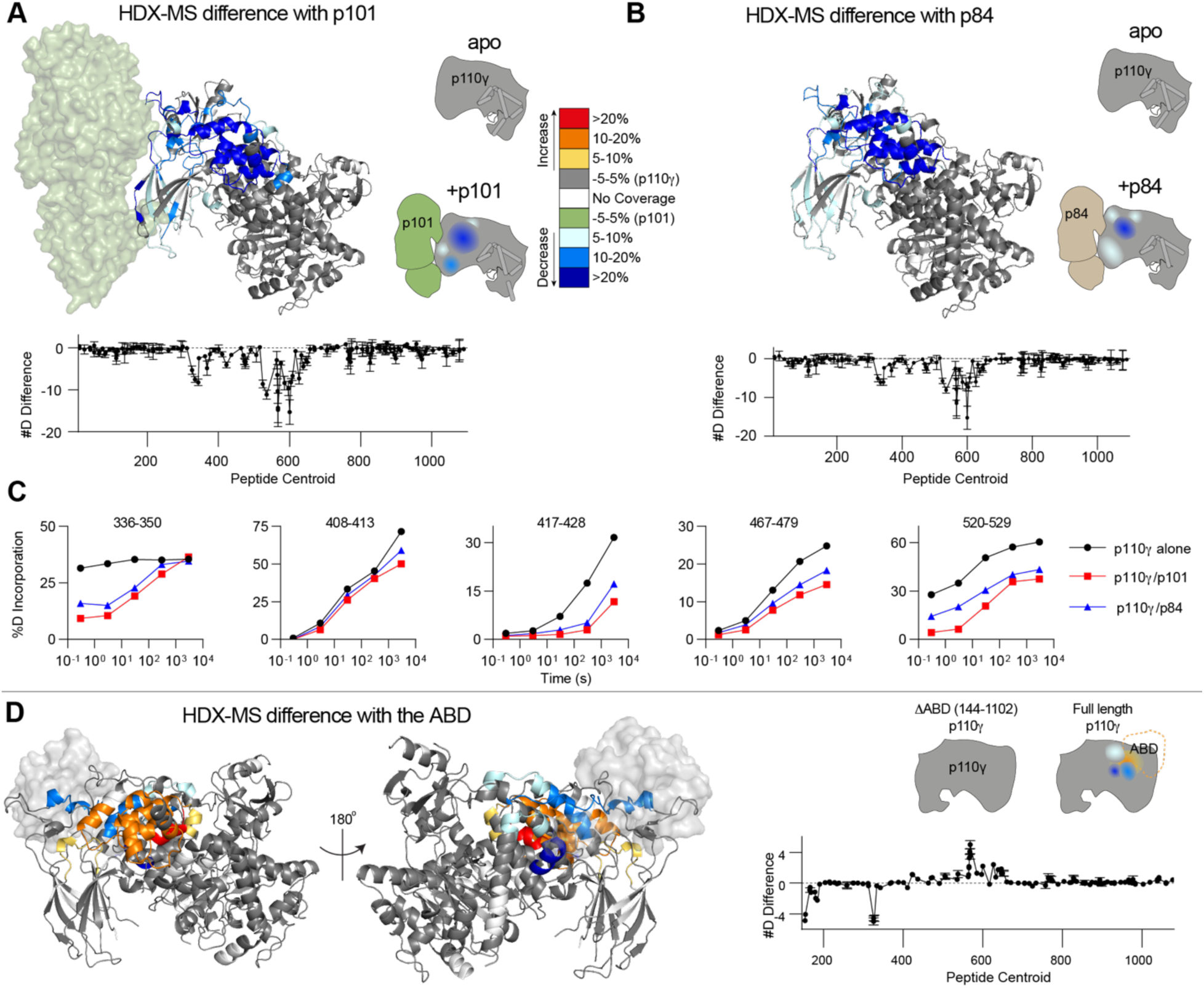
HDX-MS differences in p110γ with regulatory subunits and the ABD. **A.** HDX-MS differences in p110γ with the addition of the p101 subunit. **(A+B+D)** Peptides showing significant deuterium exchange differences (>5%,>0.4 kDa and p<0.01 in an unpaired two-tailed t-test) between conditions are colored on a cartoon model of p110γ-p101 or p110γ alone. A cartoon schematic is shown indicating the two conditions compared using HDX-MS. The number of deuteron difference for p110γ-p101 for all peptides analysed over the entire deuterium exchange time course is shown for p110γ. Every point represents the central residue of an individual peptide. Error is shown as standard deviation (n = 3). **B.** HDX-MS differences in p110γ with the addition of the p84 subunit mapped on a model of p110γ. The number of deuteron difference for p110γ-p84 for all peptides analysed over the entire deuterium exchange time course is shown for p110γ. **C.** Selected p110γ peptides that showed decreases and increases in exchange between p110γ alone, p110γ-p101, and p110γ-p84. The HDExaminer output data and the full list of all peptides and their deuterium incorporation is shown in the source data file. **D.** HDX-MS differences in p110γ with the presence of the ABD mapped on a model of p110γ. The number of deuteron difference for the p110γ ABD deletion for all peptides analysed over the entire deuterium exchange time course is shown for p110γ.

**Fig S7.**
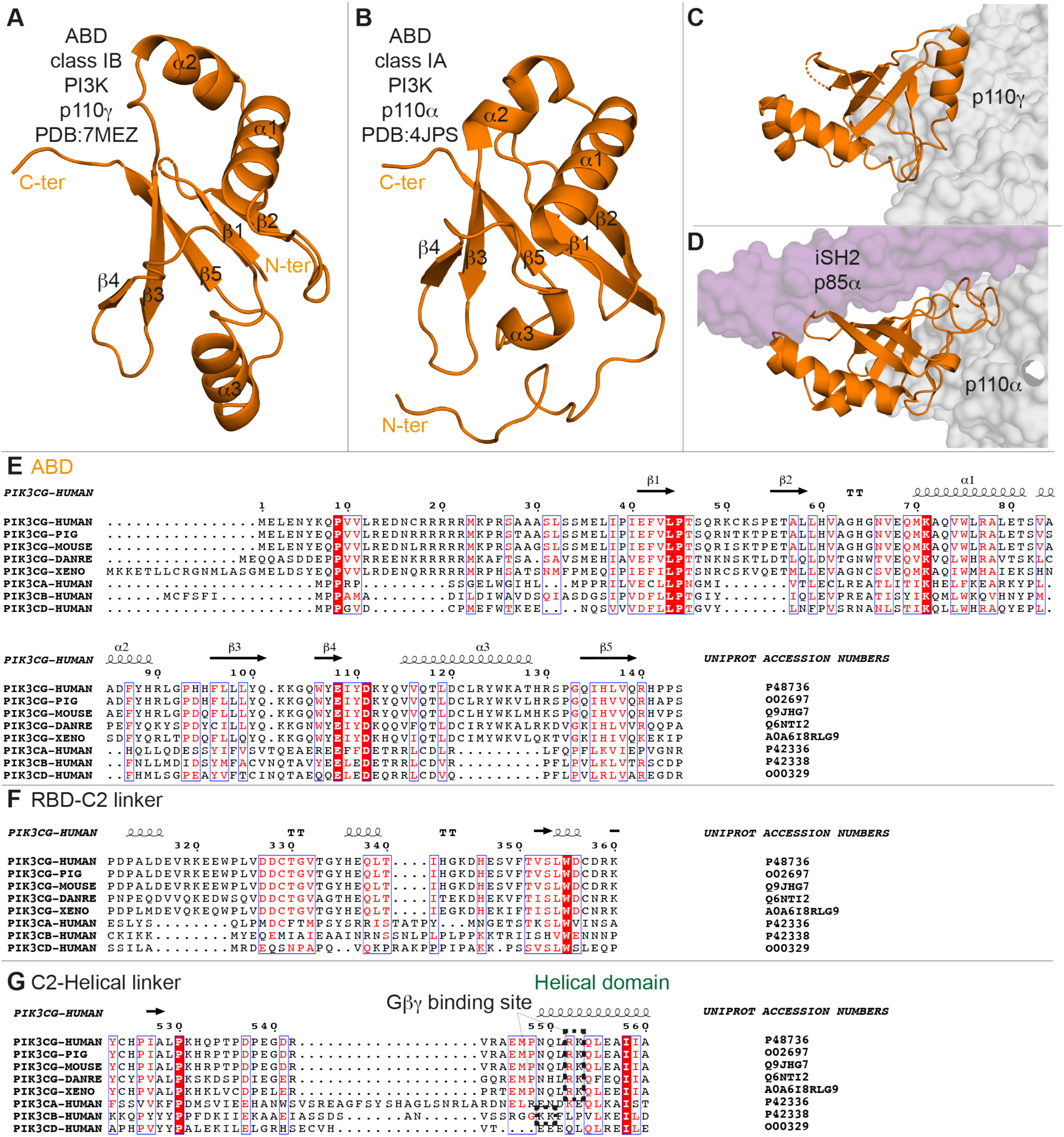
Key structural differences between p110γ and Class IA p110s. **A.** Structural model of p110γ ABD with secondary structure elements **B.** Structural model of p110α ABD with secondary structure elements **C.** Contacts made by p110γ ABD (orange) with the rest of p110γ (grey surface) **D.** Contacts made by p110α ABD (orange) with the rest of p110α (grey surface) and the p85α iSH2 domain (purple surface) **E.** Alignment showing evolutionary conservation of residues in the ABD of human p110γ with corresponding p110γ sequences from pig, mouse, Xenopus and zebrafish and corresponding class IA p110 sequences from human. The secondary structure elements of human p110γ are shown above the alignment. **F.** Alignment showing evolutionary conservation of residues in the RBD-C2 of human p110γ with corresponding p110γ sequences from pig, mouse, Xenopus and zebrafish and corresponding class IA p110 sequences from human. The secondary structure elements of human p110γ are shown above the alignment. **G.** Alignment showing evolutionary conservation of residues in the C2-helical linker of human p110γ with corresponding p110γ sequences from pig, mouse, Xenopus and zebrafish and corresponding class IA p110 sequences from human. The secondary structure elements of human p110γ are shown above the alignment. All alignments generated using ESPript 3.0.

**Fig S8.**
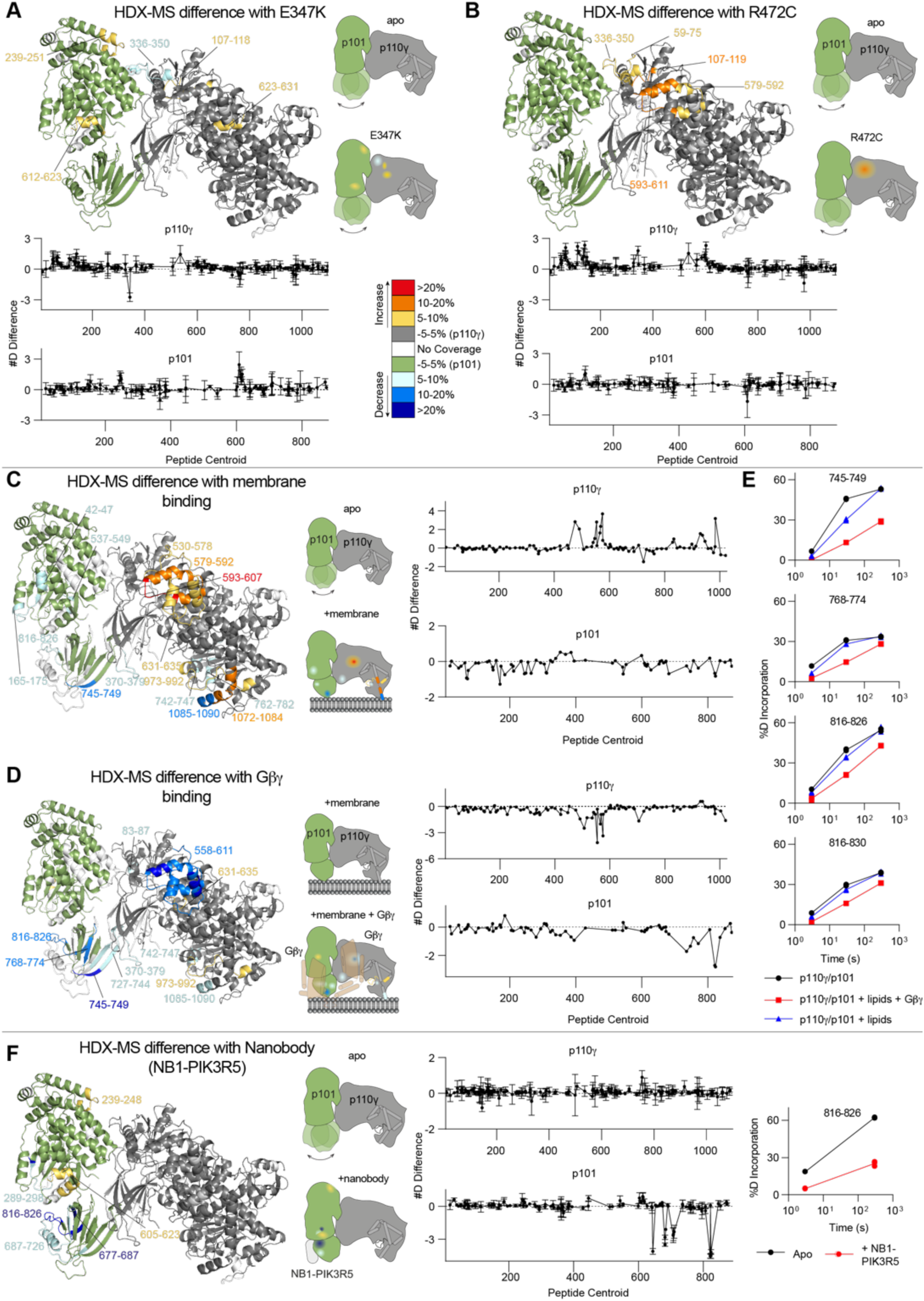
HDX-MS differences of p110γ oncogenic mutants and p110γ binding to lipids, Gβγ, and nanobody. **A.** HDX-MS differences in p110γ-p101 between wild-type p110γ-p101 and E347K. For panels **A-D + F,** peptides showing significant deuterium exchange differences (>5%,>0.4 kDa and p<0.01 in an unpaired two-tailed t-test) between conditions are colored on a cartoon model of p110γ-p101. A cartoon schematic is shown indicating the two conditions compared using HDX-MS. The number of deuteron difference for E357K for all peptides analysed over the entire deuterium exchange time course is shown for p110γ and p101. For all #D graphs, every point represents the central residue of an individual peptide, with error shown as standard deviation (n = 3). **B.** HDX-MS differences in p110γ-p101 between wild-type p110γ-p101 and R472C mapped on a model of p110γ-p101. The number of deuteron difference for R472C for all peptides over the full time course of exchange is shown for p110γ and p101. **C.** HDX-MS differences in p110γ-p101 upon binding to membrane mapped on a model of p110γ-p101. The number of deuteron difference for all peptides analysed over the full time course of exchange is shown for p110γ and p101. **D.** HDX-MS differences in p110γ-p101 upon binding to Gβγ mapped on a model of p110γ-p101. The number of deuteron difference for all peptides analysed over the full time course of exchange is shown for p110γ and p101. **E.** Selected p101 peptides that showed decreases and increases in exchange between p110γ-p101 alone, p110γ-p101 with membrane, and p110γ-p101 with membrane and Gβγ. The individual data points are shown on the graph (n=2). The HDExaminer output data and the full list of all peptides and their deuterium incorporation is shown in the source data file. **F.** HDX-MS differences in p110γ-p101 bound to NB1-PIK3R5 mapped on a model of p110γ-p101. The number of deuteron difference for all peptides analysed over the entire deuterium exchange time course is shown for p110γ and p101. A single p101 peptide showing H/D exchange data is shown. The individual data points are shown on the graph (n=3).

**Fig S9.**
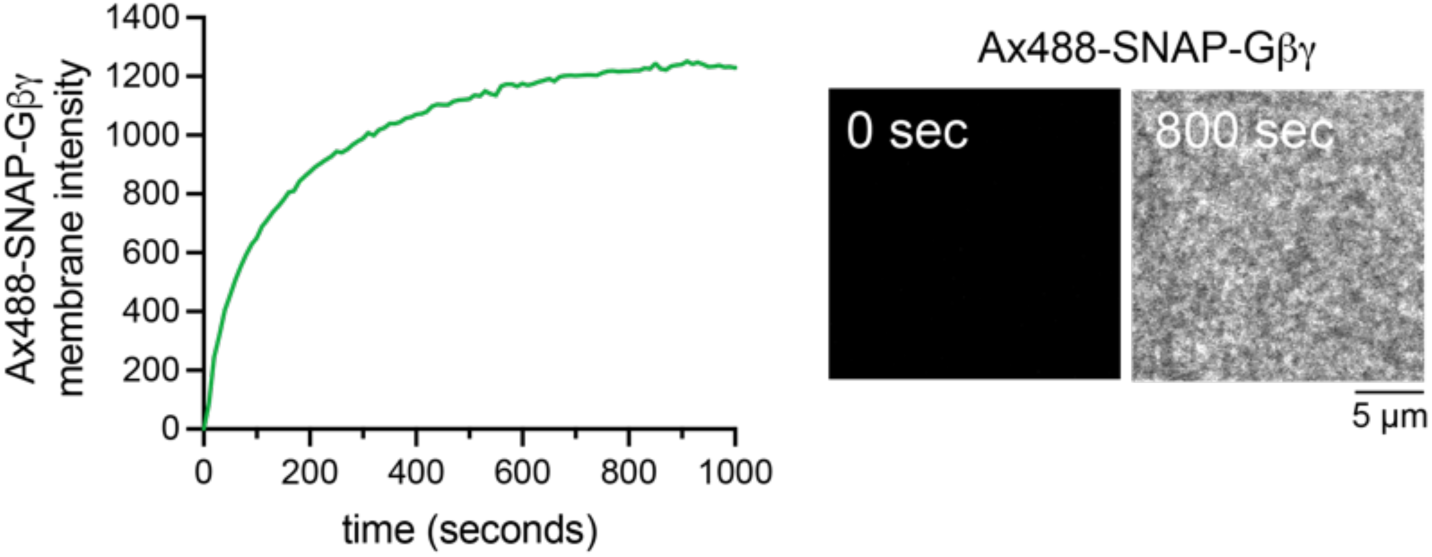
Absorption of lipidated Gβγ on membrane surface. Trace showing absorption kinetics of 400 nM Gβγ (0.5% Alexa488-SNAP-Gβγ) on a supported membrane. Representative TIRF-M images of Alexa488-SNAP-Gβγ membrane localization at the start and end of the absorption profile are shown.

**Table S1.**
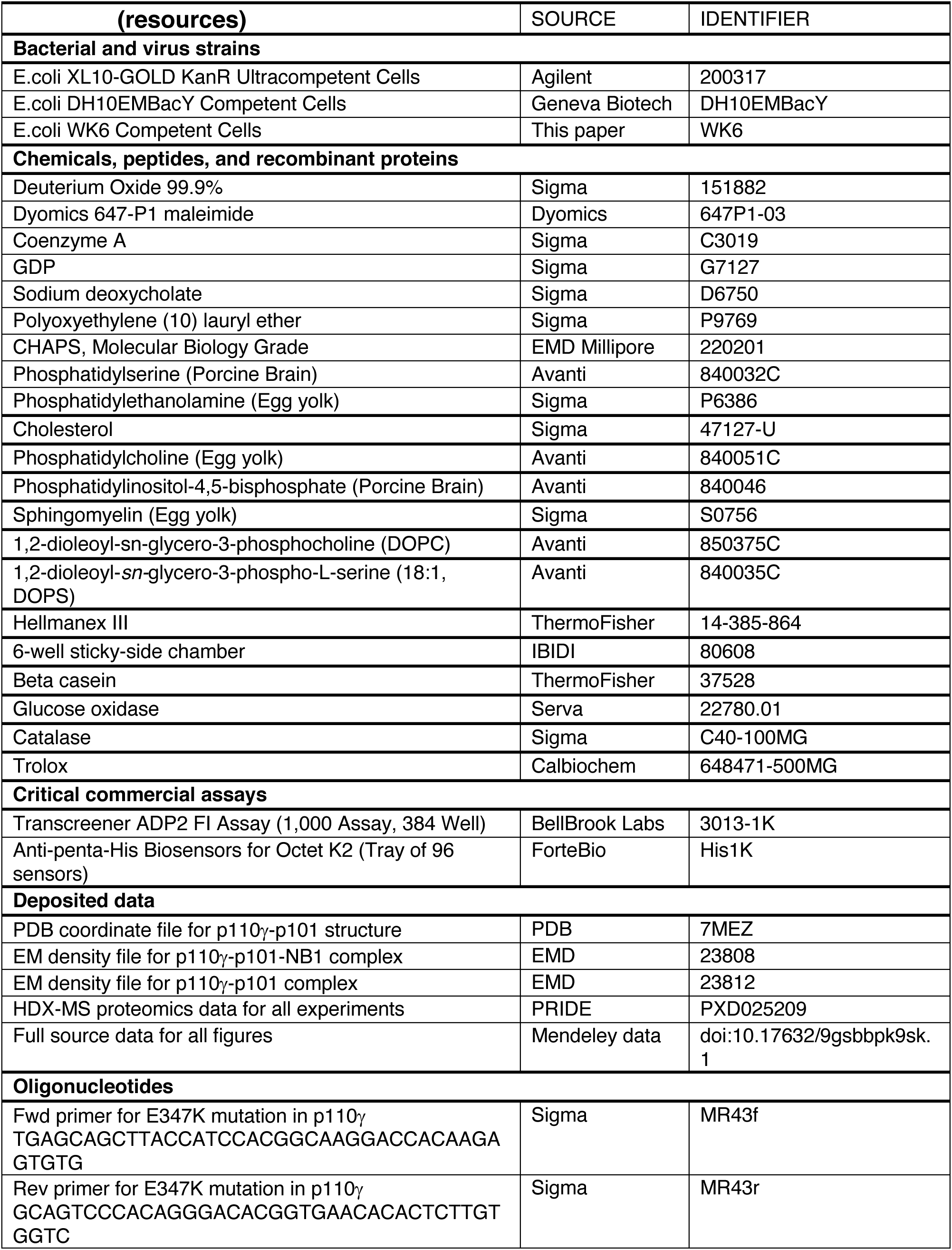
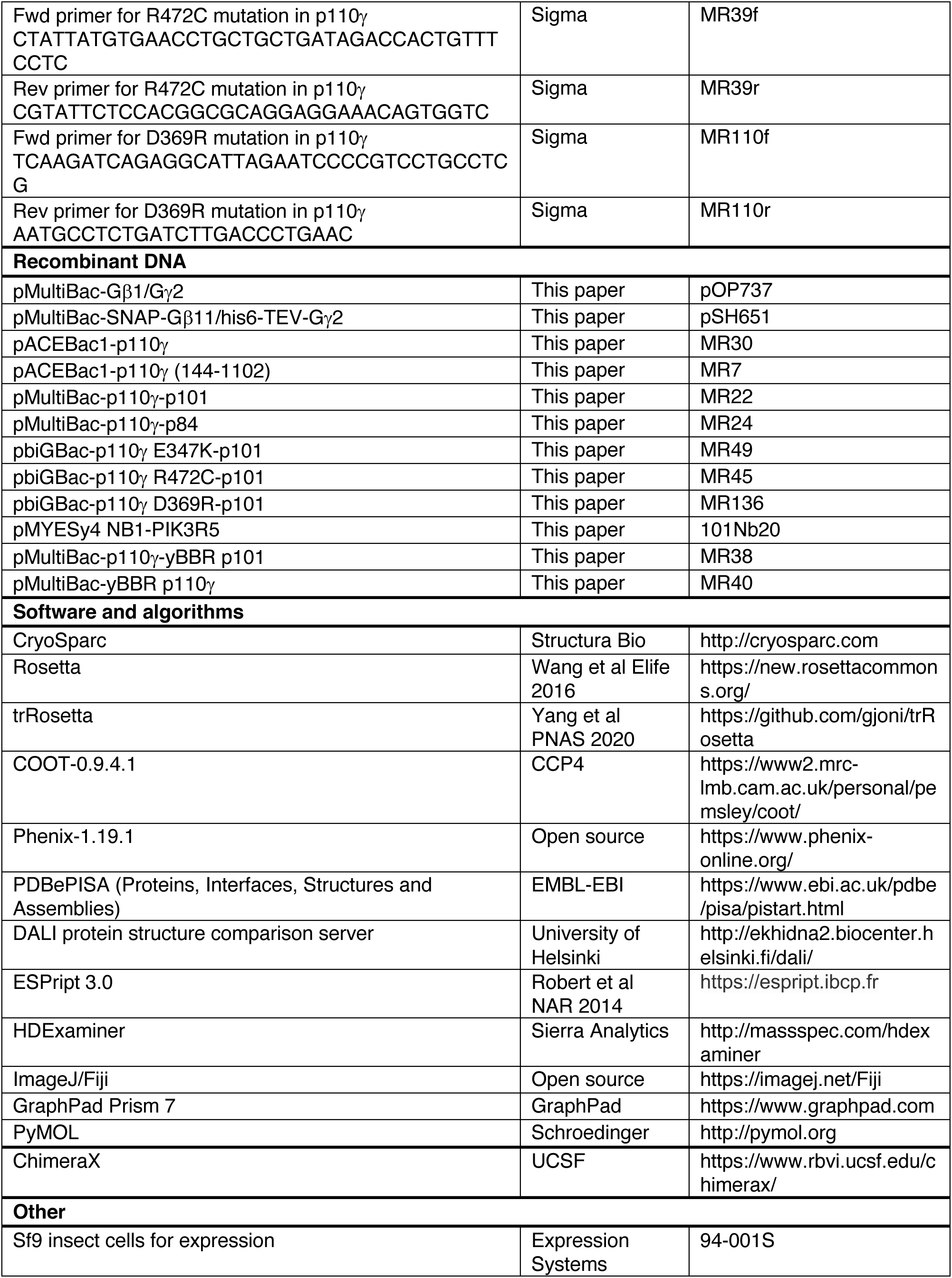

**Table S2.**
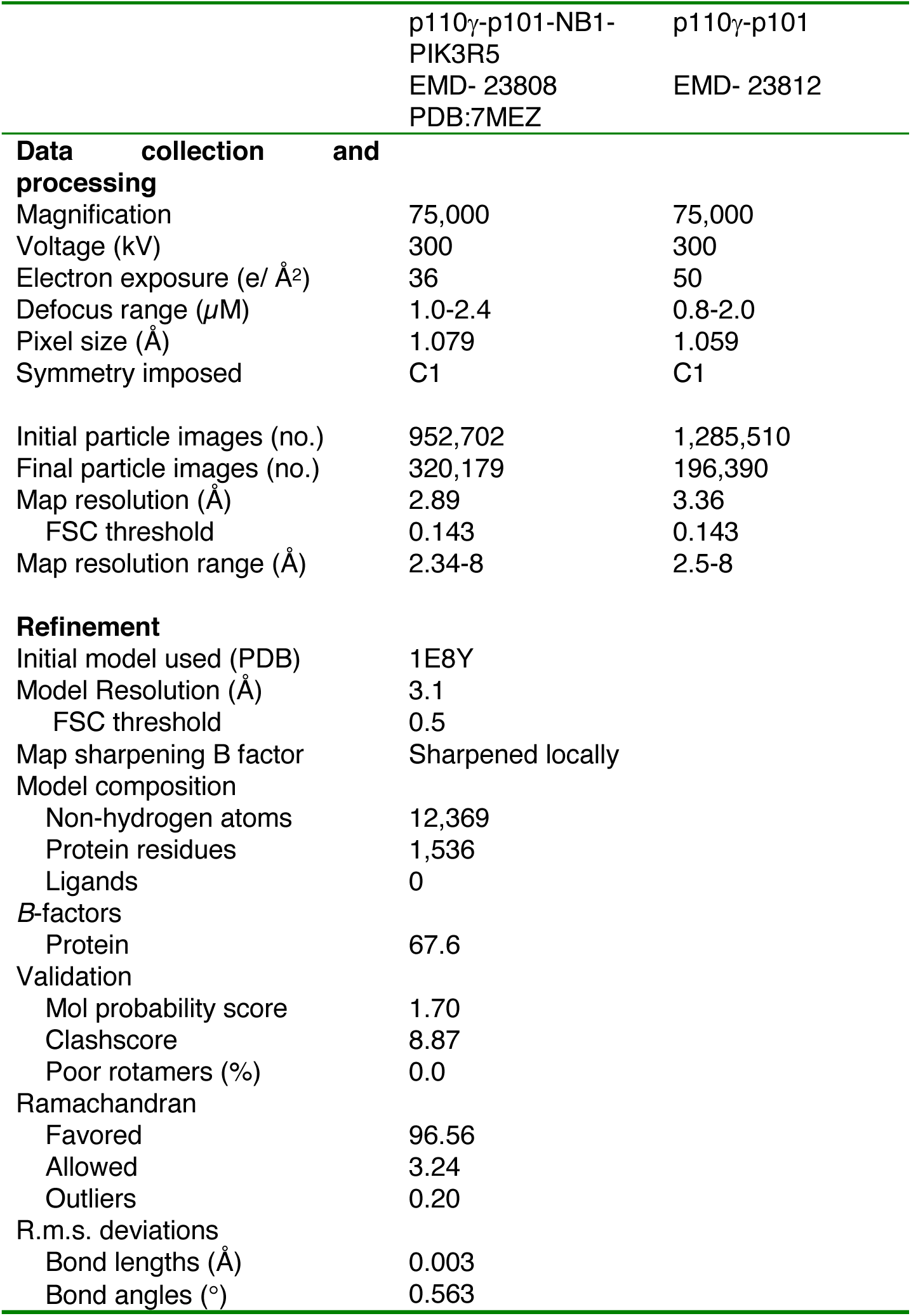
Cryo-EM data collection, refinement, and validation statistics

**Table S3.** HDX-MS data analysis table

| Data set | Apo p110γ | p110γ/p101 | p110γ/p84 | Apo p110γ/p101 | p110γ/p101 + nanobody |
| --- | --- | --- | --- | --- | --- |
| HDX reaction details | %D<sub>2</sub>O=91.7%<br>pH<sub>(read)</sub>=7.5<br>Temp=18°C | %D<sub>2</sub>O=91.7%<br>pH<sub>(read)</sub>=7.5<br>Temp=18°C | %D<sub>2</sub>O=91.7%<br>pH<sub>(read)</sub>=7.5<br>Temp=18°C | %D<sub>2</sub>O=86.8%<br>pH<sub>(read)</sub>=7.5<br>Temp=18°C | %D<sub>2</sub>O=86.8%<br>pH<sub>(read)</sub>=7.5<br>Temp=18°C |
| HDX time course (seconds) | 3, 30, 300, 3000 | 3, 30, 300, 3000 | 3, 30, 300, 3000 | 3, 300 | 3, 300 |
| HDX controls | N/A | N/A | N/A | N/A | N/A |
| Back-exchange | Corrected based on %D<sub>2</sub>O | Corrected based on %D<sub>2</sub>O | Corrected based on %D<sub>2</sub>O | Corrected based on %D<sub>2</sub>O | Corrected based on %D<sub>2</sub>O |
| Number of peptides | 165 | 165 | 165 | p110γ: 181<br>p101: 130 | p110γ: 181<br>p101: 130 |
| Sequence coverage | 93.4% | 93.4% | 93.4% | p110γ: 89.6%<br>p101: 80.6% | p110γ: 89.6%<br>p101: 80.6% |
| Average peptide /redundancy | Length=12.9<br>Redundancy= 1.9 | Length= 14.4<br>Redundancy= 1.9 | Length= 14.4<br>Redundancy= 1.9 | p110γ: Length= 14.2<br>Redundancy= 2.3<br>p101: Length= 13.9<br>Redundancy= 2.1 | p110γ: Length= 14.2<br>Redundancy= 2.3<br>p101: Length= 13.9<br>Redundancy= 2.1 |
| Replicates | 3 | 3 | 3 (2 300s, 2 3000s) | 3 | 3 |
| Repeatability | Average StDev=0.6% | Average StDev=0.6% | Average StDev=0.6% | Average StDev<br>p110γ=0.5%<br>p101=0.7% | Average StDev<br>p110γ=0.7%<br>p101=0.8% |
| Significant differences in HDX | >5% and >0.4 Da and unpaired t-test ≤0.01 | >5% and >0.4 Da and unpaired t-test ≤0.01 | >5% and >0.4 Da and unpaired t-test ≤0.01 | >5% and >0.4 Da and unpaired t-test ≤0.01 | >5% and >0.4 Da and unpaired t-test ≤0.01 |

| Data set | Apo p110γ | p110γ ABD deletion | Apo p110γ/p101 | p110γ/p101 R472C | p110γ/p101 E347K |
| --- | --- | --- | --- | --- | --- |
| HDX reaction details | %D<sub>2</sub>O=92.0%<br>pH<sub>(read)</sub>=7.5<br>Temp=18°C | %D<sub>2</sub>O=92.0%<br>pH<sub>(read)</sub>=7.5<br>Temp=18°C | %D<sub>2</sub>O=90.5%<br>pH<sub>(read)</sub>=7.5<br>Temp=18°C | %D<sub>2</sub>O=90.5%<br>pH<sub>(read)</sub>=7.5<br>Temp=18°C | %D<sub>2</sub>O=90.5%<br>pH<sub>(read)</sub>=7.5<br>Temp=18°C |
| HDX time course (seconds) | 3, 30, 300, 3000 | 3, 30, 300, 3000 | 0.3, 3, 30, 300, 3000 | 0.3, 3, 30, 300, 3000 | 0.3, 3, 30, 300, 3000 |
| HDX controls | N/A | N/A | N/A | N/A | N/A |
| Back-exchange | Corrected based on %D<sub>2</sub>O | Corrected based on %D<sub>2</sub>O | Corrected based on %D<sub>2</sub>O | Corrected based on %D<sub>2</sub>O | Corrected based on %D<sub>2</sub>O |
| Number of peptides | 150 | 150 | p110γ: 194<br>p101: 122 | p110γ: 194<br>p101: 122 | p110γ: 194<br>p101: 122 |
| Sequence coverage | 91.0% | 91.0% | p110γ: 92.1%<br>p101: 87.5% | p110γ: 92.1%<br>p101: 87.5% | p110γ: 92.1%<br>p101: 87.5% |
| Average peptide /redundancy | Length= 14.4<br>Redundancy= 2.0 | Length= 14.4<br>Redundancy= 2.0 | p110γ: Length= 14.3<br>Redundancy= 2.5<br>p101: Length= 14.0<br>Redundancy= 1.9 | p110γ: Length= 14.3<br>Redundancy= 2.5<br>p101: Length= 14.0<br>Redundancy= 1.9 | p110γ: Length= 14.3<br>Redundancy= 2.5<br>p101: Length= 14.0<br>Redundancy= 1.9 |
| Replicates | 3 | 3 | 3 | 3 | 3 |
| Repeatability | Average StDev=0.4% | Average StDev=0.4% | Average StDev<br>p110γ=0.6%<br>p101=0.9% | Average StDev<br>p110γ=0.6%<br>p101=0.8% | Average StDev<br>p110γ=0.6%<br>p101=0.9% |
| Significant differences in HDX | >5% and >0.4 Da and unpaired t-test ≤0.01 | >5% and >0.4 Da and unpaired t-test ≤0.01 | >4% and >0.4 Da and unpaired t-test ≤0.01 | >4% and >0.4 Da and unpaired t-test ≤0.01 | >4% and >0.4 Da and unpaired t-test ≤0.01 |

**Table S4.**
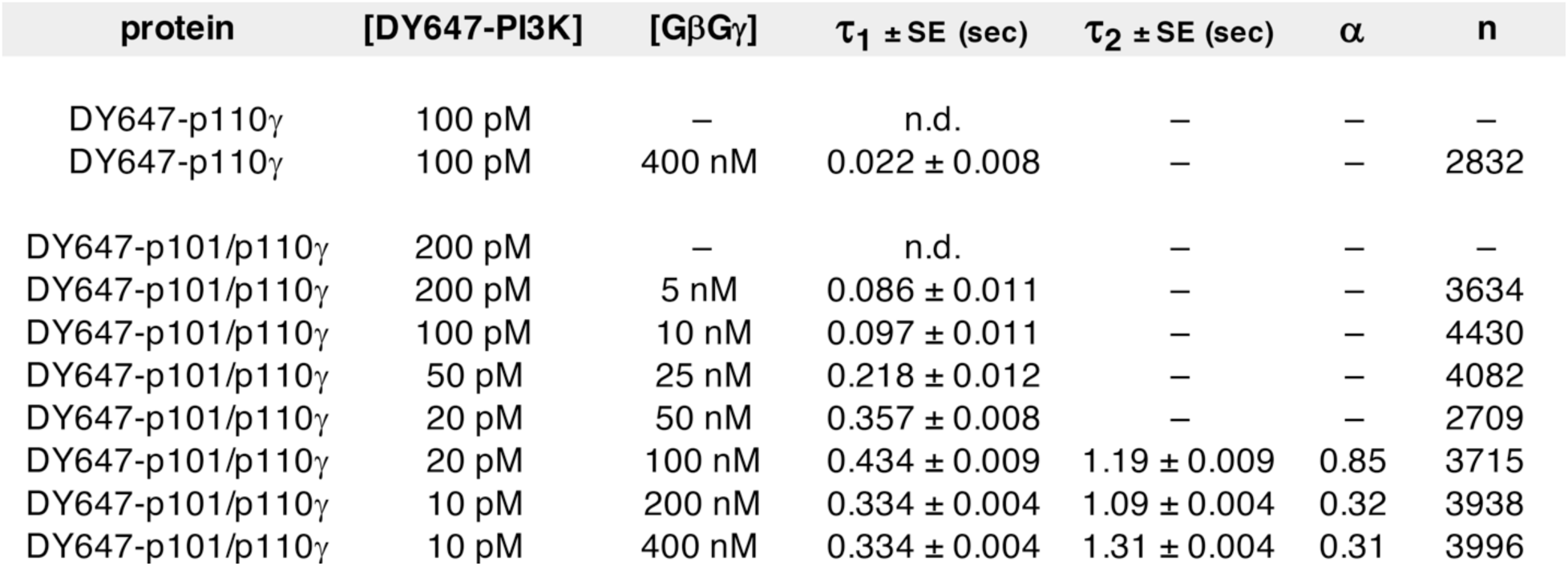
Single molecule dwell time data from TIRF-M experiments. The indicated concentration of either DY647-p101/p110γ and DY647-p110γ were flowed into the SLB containing sample chambers with varying concentrations of GβGγ (0.5% Alexa488-SNAP-GβGγ). The total number of single molecule binding events (n) reported is from 2-3 experiments. Alpha (α) is the fraction of molecules in the distribution with the short dwell time (τ_1_). Dwell time distributions were fit with either a single or double exponential decay curves. SE is the standard error from non-linear regression. Membrane composition: 95% DOPC, 5% DOPS.

